# A Flexible Quadruple-Stranded Helicate Demonstrates a Strong Binding Preference for DNA Three-Way Junctions by Induced-Fit

**DOI:** 10.1101/2025.08.21.671622

**Authors:** Hugo D. Williams, Samuel J. Dettmer, Sumit Bajpai, Michael J. Hannon

## Abstract

Nucleic acid junctions are key to many biological functions from recombination and repair to viral NA insertion, and are an attractive, functional biomolecular target. We describe a quadruple-stranded di-platinum helicate that binds both three-way (3WJ) and four-way DNA junctions (4WJ). This allows us to probe the relative importance of size and shape in junction-binder design. Despite the helicate’s tetragonal symmetry/shape being compatible with the 4WJ, microscale thermophoresis (MST), isothermal calorimetry (ITC) and gel electrophoresis competition experiments demonstrate that this metallo-supramolecule displays a stronger affinity for 3WJs (K_d_ = 12 nM) than for 4WJs (K_d_ > 4 µM) and other DNA structures. The experimental findings are supported by molecular dynamics simulations which reveal the critical role of size. Whilst the open form of the 4WJ is promoted when the helicate is in the cavity, the helicate’s small size means it is unable to maintain π contacts with all four junction base-pairs simultaneously. Although the helicate is slightly too large for the smaller 3WJ cavity, simulations and experiments show that it can open up the cavity (increasing the junction’s hydrodynamic radius) by disrupting a base-pair. The flexible helicate also responds to the cavity upon binding by favouring one enantiomer and allowing the helicate to adopt a stable final structure inside the 3WJ that is an induced-fit of the two dynamic structures (supramolecule and DNA). This contrasts with previous lock-and-key examples of junction recognition and opens up new possibilities for how to design DNA and RNA junction-binding compounds.

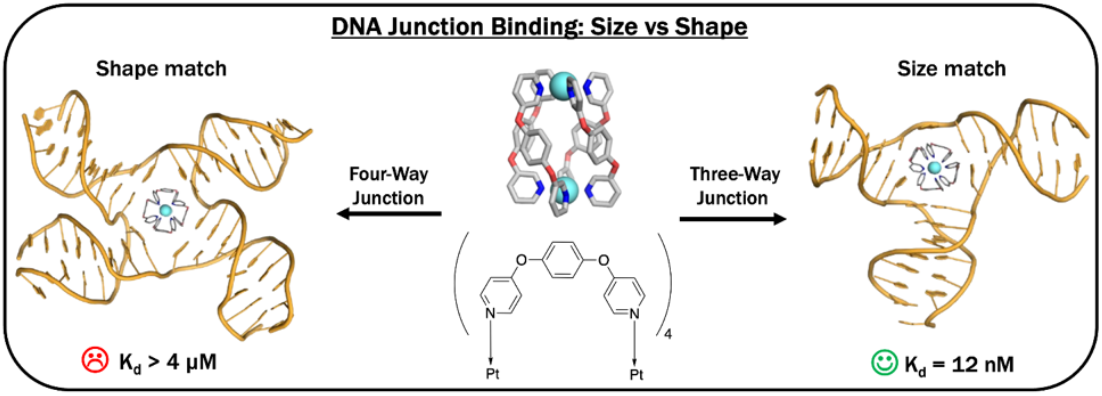

## INTRODUCTION

DNA is an important and abundant biomolecule that is the fundamental store of the genetic information governing the production of proteins and the function of cells in most organisms. DNA resides for the majority of time (and when not being processed) in a B-double helix form. Some cancer chemotherapies target DNA in this form, with typical examples being cross-linking agents, such as cisplatin and its derivatives, and intercalators, such as anthracyclines. However, given that almost all cells contain DNA in this form, these compounds lack specificity and exhibit unwanted side effects.^1–5^ Processing the encoded genetic information requires unwinding of the helix, which gives rise to the formation of non-canonical intermediate structures that could be more interesting/effective as therapeutic targets.^6^

A higher order DNA structure commonly observed in cell processes is the junction. The four-way Holliday junction (4WJ) was first proposed in 1964 as the major intermediate in homologous recombination and is understood to be an important structure in rescuing stalled replication forks and in double-strand break repair.^7–10^ It consists of four duplex domains that converge at a branchpoint, and can adopt two primary conformations, open cruciform and closed X-stacked, of which the former is most frequently observed in biological processes.^11-13^ Less common is the three-way junction (3WJ), which does not have a prominent role in normal cell processes, though it is implicated in DNA transactional processes and has been shown to form in genomic regions containing triplet nucleotide repeats. These repeats are associated with expansion disorders such as Fragile X syndrome and Huntington’s disease.^14–17^ Analogous to the 4WJ, the 3WJ comprises three duplex domains that converge at a branchpoint cavity.

Recently there has been an upsurge in interest in junctions as biological targets driven by and enabled by several supramolecular or peptidic compounds that have been identified as DNA junction binders. Cationic [M_2_L_3_]^4+^ triple-helicate metallo-supramolecular cylinders possess the right size and shape to thread beautifully through the central cavity of the 3WJ and stabilise it (Fig. 1A) as revealed crystallographically.^18^ Importantly, the binding features exquisite π–π stacking of the cylinder’s outward-facing aromatic surfaces and the branchpoint nucleobases. Organic azacryptands also bear comparable π-surfaces, allowing them to bind 3WJs in a similar way.^19^ Other helicates that do not display aromatic faces on their surface, but with similarly compatible shape–size profiles are also proposed to bind in the 3WJ cavity.^20–22^

**Figure 1.**
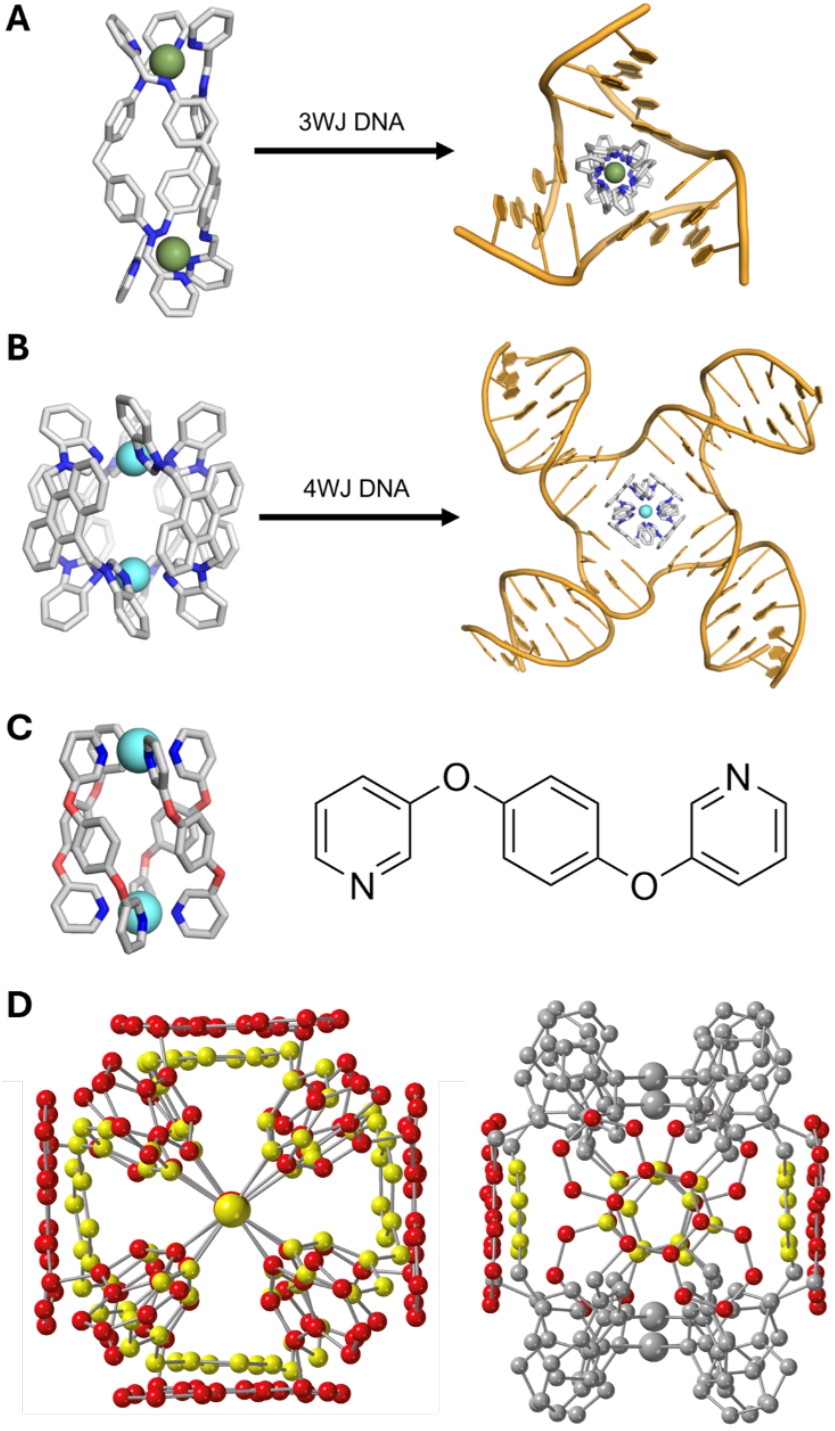
A) Iron triple helicate cylinder threading through the cavity of a hexamer 3WJ (crystal structure; PDB 2ET0).^18^ Hydrogens omitted for clarity. B) Platinum metallo-cage threading into the cavity of a DNA 4WJ (MD snapshot).^27^ Hydrogens omitted for clarity. C) Crystal structure of the Steel-McMorran palladium helicate (left) and the structure of the ligand L1 (right).^28^ D) End-on (left) and side-on (right) view of a superposition of the Pt-BIMA metallo-cage (red) and the Steel–McMorran Pd helicate (yellow). In the side-on view, only the aromatic surface carbons are coloured for clarity. The Steel–McMorran helicate exhibits a smaller radius than Pt-BIMA, as well as displaying a smaller π surface.

Work on 4WJ binders has often focussed on small (intercalator-style) molecules which target the closed form of the junction,^23, 24^ or small peptides that bind, possibly as a dimer, to a partially-open form (though the nature of their molecular interactions with the 4WJ remain elusive).^25^ We recently reported that organometallic pillarplexes bearing imidazole surfaces and a square symmetry are able to bind and stabilise the open conformation.^26^ Taking inspiration from [M_2_L_3_]^4+^ 3WJ binders, we have further shown that a rigid [M_2_L_4_]^4+^ metallo-cage presenting anthracene ligand surfaces (Pt-BIMA) binds elegantly inside the 4WJ open cavity, wherein the anthracene surfaces stack optimally with the four branchpoint base pairs (Fig. 1B).^27^

Yet despite these successes, the detailed understanding of how to bind DNA junction structures remains in its infancy. To unlock and fully exploit the exciting biological potential of such binding, more detailed knowledge is needed. In particular, we note that despite their 4WJ preference and ideal symmetry for 4WJ binding, both the pillarplex and the Pt-BIMA (anthracene lined) metallo-cage also bind 3WJs, albeit with lower affinity. This is facilitated by the opening of a branchpoint base pair, which expands the 3WJ cavity to accommodate the larger compounds. Opening up of DNA junctions is interesting and exciting *per se*, but it also hints at: (i) a tension between size match and shape/symmetry match of the binding agent with the junction cavity, and (ii) the dynamic nature of these DNA structures and their potential for changes in conformation in response to the binder. To start to explore this in greater detail, we now describe the junction binding of quadruple-stranded metallo-cages that are smaller than the previous anthracene cages which allows us to probe the relative importance of size versus symmetry in junction binding.

## RESULTS AND DISCUSSION

The first coordinatively-saturated quadruple-stranded helicate was described by Steel and McMorran assembled from four bis-pyridyl ligands (L1) with two square planar palladium(II) centres.^28^ Their original pioneering work showed that the resulting helicate/cage could encapsulate a hexafluorophosphate anion and more recently they have shown that different encapsulated anions affect the dimensions of the structure by changing the helical pitch, giving possible Pd–Pd distances of between 7.4 and 8.8 Å.^29^ Inspired by this work, many other examples of palladium(II) based quadruple-stranded Pd_2_L_4_ complexes have been described in the literature, with a particular focus on how their internal cavities can be used to capture, deliver and release cargo in response to stimuli.^30–40^ The original Steel and McMorran palladium(II) helicate possesses a number of properties which make it an attractive complex for DNA junction binding: it is cationic, has dimensions similar to those of other junction-binders, and displays four outward facing aromatic surfaces (Fig. 1C). Compared to the 4WJ-binding anthracene Pt-BIMA cage, this helicate is slightly smaller (Fig. 1D), contains only one aromatic ring per surface and is much more flexible (given its ability to dynamically respond to encapsulated anions). Thus, it is an ideal test system for us to start to probe the effects of size and symmetry in junction-binder design.

### Synthesis and Characterisation of the Compound

The ligand (L1) was synthesised as reported previously by the condensation of dibromobenzene with 3-hydroxypyridine.^41^ We then prepared the previously reported palladium helicate [Pd_2_(L1)_4_](BF_4_)_4_ by reaction with [Pd(MeCN)_4_](BF_4_)_2_ in acetonitrile,^28^ however we found this metallo-cage to have poor stability in aqueous Tris buffer (Fig. S10), which we commonly utilise for biophysical experiments. To address this stability issue and enable effective study of the system, we decided to instead prepare a new complex, a platinum(II) analogue [Pt_2_(L1)_4_](NO_3_)_4_. While palladium(II) metallo-cages are well studied, platinum(II) analogues are much less well established.^27, 42–44^

Reaction of the L1 ligand with [Pt_2_(DMSO)_2_Cl_2_] in acetonitrile under reflux for 48 hours led to a mixture of 4-ligand M_2_L_4_ and 5-ligand M_2_L_5_ species which were difficult to separate. Using DMSO as solvent and warming at 120 ^°^ C led to a similar initial mixture, but over a period of 24 hours the mixture rearranged into the desired [Pt_2_(L1)_4_]^4+^ helicate (Fig. 2). Severin has crystallographically characterised an example of a 5-stranded Pd_2_L_5_ species with a different bis-monodentate ligand, in which three ligands bridge both metals and the other two act as monodentates but are stacked into the helical structure.^45^ The observed 5-ligand intermediate with this ligand L1, likely has a similar structure, and is observed as a stable intermediate due to the lower kinetic lability of Pt(II).

**Figure 2.**
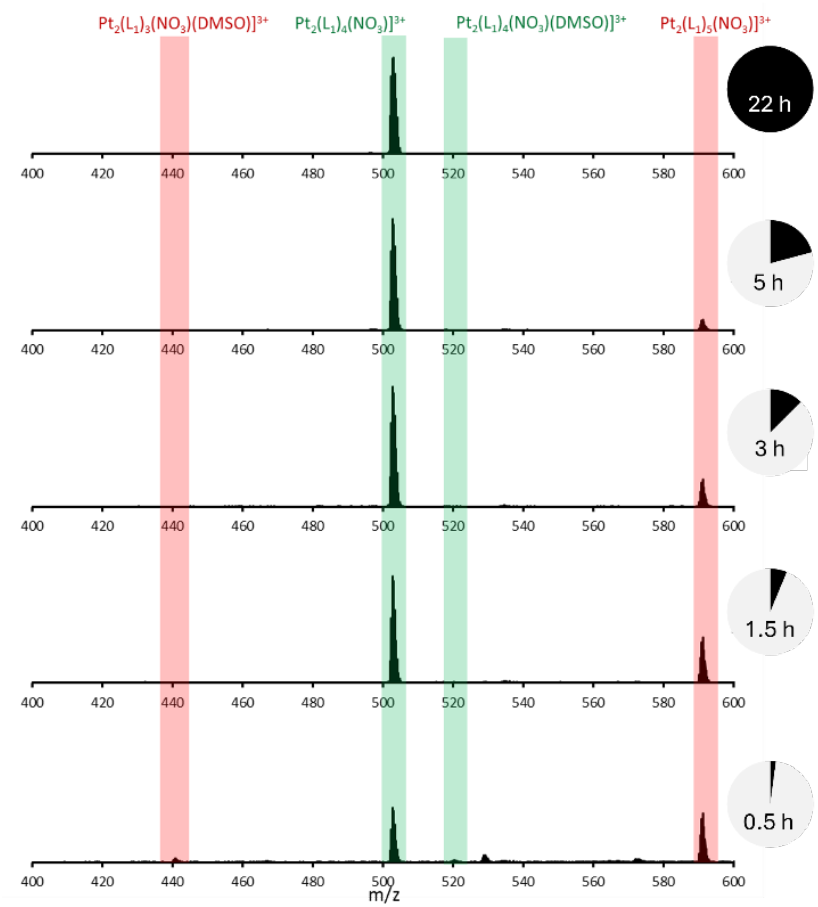
Monitoring the formation of the [Pt_2_(L1)_4_](NO_3_)_4_ complex by ESI-MS. Highlighted in green are peaks corresponding to the desired product and highlighted in red are peaks corresponding to a 5 ligand species, which decreased over time.

Though the Pt_2_(L1)_4_ complex is formally a tetra-cation, the 3+ species was the dominant species seen in ESI-MS due to efficient encapsulation of a nitrate anion in the cavity of the helicate (this reflects the anion encapsulation seen in the palladium(II) analogue).^28^ Formation of the pure platinum Pt_2_(L1)_4_(NO_3_)_4_ (Pt helicate) complex as a discrete species (no penta-ligand species) was confirmed by ^1^H NMR spectroscopy and elemental analysis. The Pt helicate showed a much improved solution stability compared to the Pd helicate in aqueous buffer conditions, shown by UV-Vis spectroscopy (Fig. S11). Though Steel and McMorran’s crystal structure presents a helical compound with axial chirality, we observe for the Pt complex (as they do for their Pd complex) only one phenyl proton signal in the NMR spectrum. Steel and McMorran postulate that this is due to the rapid interconversion of the helicate between the *M* and *P* forms in solution,^29^ though this NMR feature may also be explained by spinning of the phenyl ring.

### Screening Three-Way and Four-Way Junction Binding

Polyacrylamide gel electrophoresis (PAGE) was first used to qualitatively assess the junction binding of the compound (Fig. 3A). To investigate the 4WJ, four individual oligomers, which collectively form a 4WJ were mixed with increasing concentrations of the Pt helicate in buffer (1x TB, 10 mM NaCl). A gel shift is observed in the slowest-moving band (corresponding to the 4WJ), indicating that the complex binds the 4WJ structure (Fig. 3A, lanes 13–19). Additionally, two strong intermediate bands appear between the single-strands band and the 4WJ band, corresponding to a two-stranded Y-fork and a three-stranded *pseudo*-3WJ (p3WJ), which both increase in intensity as the concentration of the Pt helicate increases. This behaviour contrasts with that of the Pt-BIMA metallo-cage which preferentially binds the 4WJ and forms these bands much more faintly,^27^ but is very similar to the behaviour of the Ni cylinder (Fig. 3A, lane 5), which is an excellent 3WJ binder.

**Figure 3.**
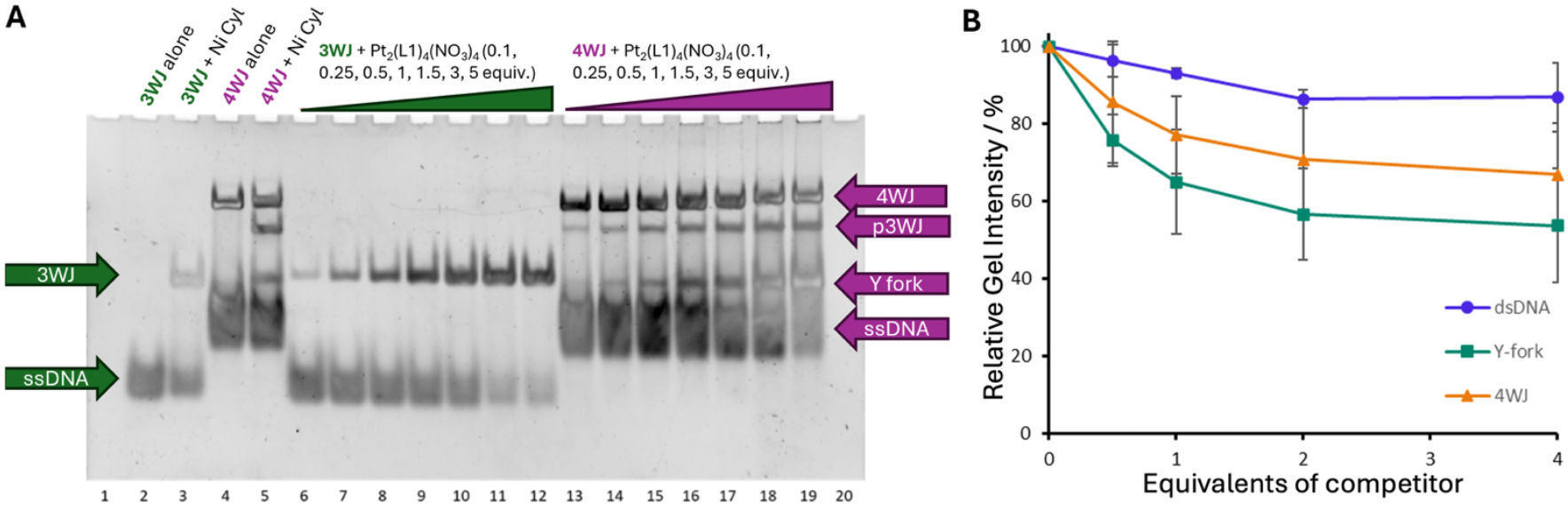
A) PAGE gel of increasing concentrations of Pt helicate incubated with DNA 3WJ and 4WJ structures. Controls contain the 3WJ and 4WJ with no junction binder (lanes 2 and 4) and the 3WJ and 4WJ with a known junction binder (Ni cylinder, lane 3 and 5 respectively). The 3WJ with 0.1, 0.25, 0.5, 1, 1.5, 3, and 5 equivalents of Pt helicate is shown in lanes 6–12. The 4WJ structure with 0.1, 0.25, 0.5, 1, 1.5, 3, and 5 equivalents of Pt_2_(L1)_4_(NO_3_)_4_ is shown in lanes 13–19. B) PAGE competition curves. The gel bands were measured and the relative 3WJ–FAM intensity is plotted as a function of the equivalent of other DNA competitors.

The 3WJ was investigated using three individual strands of a 3WJ-forming sequence that, unlike the 4WJ, do not assemble together in the absence of an appropriate binder because the 3WJ is entropically disfavoured at room temperature (Fig. 3A, lane 2). However, in the presence of a binder the 3WJ is stabilised and observed as a slower running band (Fig. 3A, lane 3). On incubation of the 3WJ oligomers with increasing concentrations of Pt helicate, the 3WJ is stabilised and the slower band appears (Fig. 3A, lanes 6–12), becoming more intense at higher concentrations of complex. The migration of the 3WJ band formed with the Pt helicate is marginally slower in comparison to the band formed with the [Ni_2_L_3_]^4+^ cylinder control, and this is particularly noticeable at lower concentrations of complex. We have previously seen this (albeit more noticeably) with the Au pillarplex and the Pt-BIMA metallo-cage, both of which are large and prefer 4WJ, but bind and open the 3WJ.^26, 27^ We attribute this slower shift to the breaking and opening of a branchpoint base pair.

Given the ability of the platinum complex to bind different DNA structures, the binding preference was established by a PAGE competition assay (Fig. S12). A fluorescently labelled 3WJ (3WJ–FAM) was mixed with 1 equivalent of Pt helicate and 0.5, 1.0, 2.0 or 4.0 equivalents of a (unlabelled) competitor DNA structure. The intensity of the 3WJ–FAM band could then be measured (Fig. 3B) without the need for staining, eliminating the difficulty of uniform staining and band overlaps. At 4 equivalents, dsDNA did not compete strongly, leading to only a ∼13% decrease in the amount of 3WJ. Y fork and 4WJ were not much more competitive, showing only a ∼22% and ∼33% decrease in the 3WJ respectively (at 4 equivalents). In contrast, the same PAGE competition experiment with the Pt-BIMA metallo-cage shows a ∼30% decrease in the 3WJ band at just 0.5 equivalents, with the signal becoming undetectable at 4 equivalents (Figs. S13–S14). This reveals that the Pt helicate displays different behaviour to Pt-BIMA and a strong preference for the 3WJ compared to dsDNA, Y-fork structures, or 4WJ. This is surprising given the four-fold symmetry of the compound.

### Molecular Dynamics and Circular Dichroism Reveal a “Hand-in-Glove” 3WJ Binding

With the PAGE results in mind, we sought to explore and understand the binding behaviour of the complex at the atomic level using molecular dynamics (MD) simulations. Starting from a reported crystal structure of the palladium compound, the Pd atoms were changed to Pt atoms and the centrally bound perchlorate anion was replaced by a nitrate anion. This was then DFT optimised and the MD coordinates and topology files generated using the MCPB workflow.^46^ Files for both formal enantiomers (*M* and *P*) were generated and both were explored as starting points for the simulations (despite the expected rapid interconversion in solution by a twisting mechanism). Simulations of the helicate alone (absence of DNA) in water and 50 mM NaCl showed that the nitrate anion could rapidly leave what is quite an open cavity. This is also observed experimentally with facile exchange of that anion in the Pd complex.^29^ The complex was thus reparameterised as a tetracation and simulations with the DNA were therefore started with the cylinder as a tetracation without internal anions, though chlorides could potentially enter during the simulations.

Starting with the 3WJ, as an initial starting position the helicate was placed directly inside the 3WJ allowing the simulation to focus on the consequences of the cavity binding. Across all simulations (3 per enantiomer, 24 µs cumulative time), the helicate was initially unable to adopt a stable binding position whilst the branchpoint bases remained paired, reflected by it rotating within the cavity – though it remained inside the cavity. Eventually, in all cases, the opening of a base pair was observed (as expected from the experimental PAGE shift) allowing the helicate to find a more stable position (Figs. 4A, 4B), in which it no longer freely rotated. In this conformation, the larger adenine base is displaced from its thymine partner, becoming unpaired and allowing the ligand of the helicate to take its place and the phenyl moiety to interact with the adjacent base pair. This positions the helicate in such a way that its other ligands can *π*-stack with the other two branchpoint base pairs (Fig.4B). The thymine in the frayed base pair was seen to remain in place and *π*-stack with one of the coordinating pyridyl groups on the cylinder – this interaction is seemingly important for locking the helicate in this conformation as it was rarely disrupted once this position was attained (Fig. S15). In 5 out of 6 simulations, the helicate remained in this position for the remainder of the simulation time. The other simulation saw the reformation of the frayed base pair, the fraying of an adjacent branchpoint base pair, and the relocation of the helicate into the new position, attaining an equivalent conformation, which was then held for the rest of the simulation (Fig. S15).

**Figure 4.**
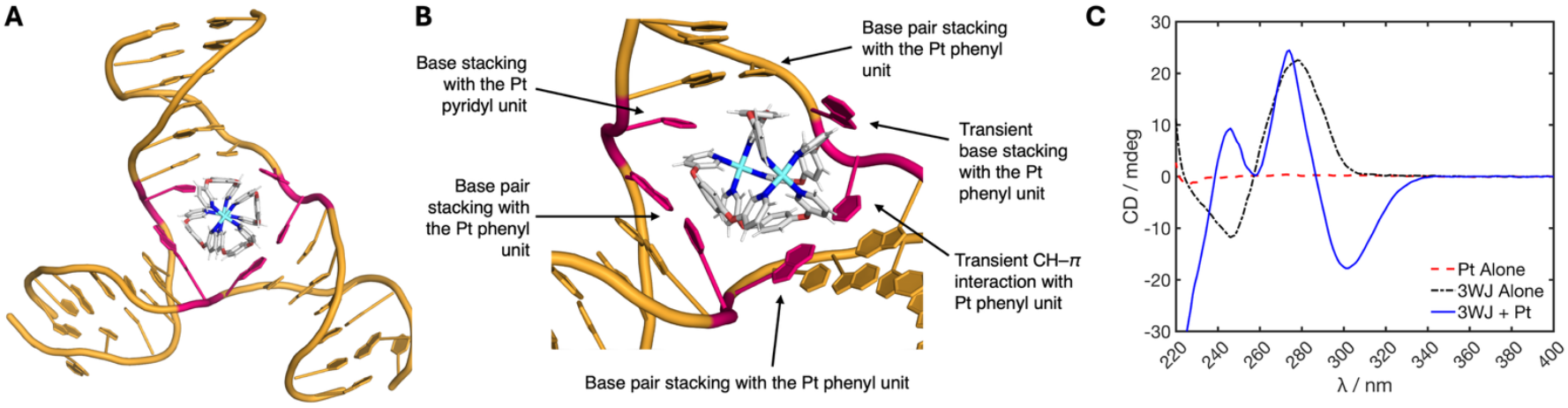
A) Representative MD snapshot of the *P* enantiomer of the Pt helicate bound inside a DNA 3WJ, showing the stable conformation adopted after the helicate displaces a base (branchpoint bases shown in pink); B) Close up of the *P* enantiomer bound inside the 3WJ cavity with labels identifying the commonly observed interactions in the simulations. Interactions labelled as transient are observed to dynamically fluctuate between bound and unbound states over the timescale of the simulations. Representative simulation videos are provided in the supplementary information. C) CD spectra of the 3WJ oligomers (5 µM, black), Pt helicate alone (5 µM, red) and 3WJ + 1 equiv. Pt helicate (5 µM, blue). All samples contained 10 mM HEPES and 100 mM NaCl.

Interestingly, in simulations starting with the *M* enantiomer, upon adopting the stable induced-fit position inside the frayed 3WJ, interconversion to the *P* enantiomer was observed in all cases indicating both that this enantiomer has the better fit and that the flexibility of this helicate is an important factor in attaining the best fit (Fig. S16). This additionally indicates that not only does the 3WJ respond to the presence of the helicate by opening a base pair, but the helicate also responds to the 3WJ cavity. One simulation starting with the *M* enantiomer captured a metastable state in which a base pair was fully opened (both bases were dislocated), the chirality of the helicate did not invert and the adenine (rather than the thymine) base was able to interact with a coordinating pyridyl unit, however this showed far more dynamic freedom in the DNA and eventually converted into the stable binding mode seen in all other simulations after ∼5.5 µs (Fig. S16). From these simulations, the interaction of the Pt helicate with the 3WJ can thus be considered a “hand-in-glove” interaction, in contrast to the “lock-and-key” interaction exhibited by the Ni cylinder.^18^

Circular dichroism (CD) was employed to assess experimentally whether one enantiomer binds to the 3WJ preferentially over the other (Fig. 4C). Enantiomers (at the same concentration) absorb circularly polarised light with equal and opposite intensities, but because the helicate exists in solution as a (dynamically interconverting) racemic mixture, it does not exhibit significant CD peaks, as the signal arising from one enantiomer cancels out the signal from the other. The Pt helicate was mixed with 1 equivalent of the 3WJ strands in buffer (10 mM HEPES and 100 mM NaCl). As seen in PAGE, the 3WJ strands do not assemble spontaneously into a 3WJ and so give a characteristic signal for single-stranded DNA with a positive peak at 280 nm and a negative peak at 245 nm. Upon binding of the Pt helicate, a strong negative induced CD band appears at 290–340 nm, and sharper positive peaks appear at 245 and 270 nm, aligning well with the compound’s UV-Vis absorbance peaks (Fig. S9). The presence of a strong induced CD signal is indicative of an excess of one enantiomer in the solution (arising by induced-fit binding to the 3WJ), consistent with the MD observations.

In further simulations, the starting position of the Pt helicate was located outside (but close to) the cavity. 18 simulations (at least 1 µs each) sampled six different starting positions on both the major and minor groove sides (Fig. S17). 16 simulations captured binding at or inside the cavity, of which 10 simulations observed the helicate quickly adopt the stable binding conformation as seen in the earlier simulations. The other 6 all saw the helicate approach from the minor groove side and bind at or partially inside the cavity, indicating that entry from the major groove side is more facile. 2 simulations did not observe entry into the cavity and instead saw the helicate bind transiently at the duplex termini. A similar end-stack binding mode is also seen for the triple-helicate cylinders both as a second mode observed in crystallography (DNA and RNA)^18, 47, 48^ and a transient binding in RNA simulations^49^; junction-binding is the dominant mode in solution.^50^

### Comparison with a “Lock-and-Key” Three-Way Junction Binder

The PAGE and MD results indicate that the Pt helicate preferentially binds the 3WJ over the 4WJ, despite its symmetrical structure. We were thus interested how the induced-fit binding of this quadruple-stranded helicate compared to the perfect fit of the triple-stranded cylinder.

To evaluate the strength of the interactions with the 3WJ, we first used isothermal titration calorimetry (ITC), which directly measures ΔH and provides the full thermodynamic profile, including the binding constant (K_d_) and stoichiometry. Titration of the Pt helicate into a pre-folded 3WJ (3WJ-T_6_), formed from a single DNA strand, yielded a 1:1 binding ratio, consistent with a junction cavity binder, and a K_d_ of 9.94 ± 4.36 × 10^−9^ M at 25 °C (Fig. S18). A similar ITC experiment with the Ni cylinder also showed 1:1 binding and a K_d_ of 7.17 ± 0.51 × 10^−9^ M (Fig. S19). This might suggest that the two compounds bind with similar affinity, however inspection of the binding curves generated from the ITC data reveals a very sharp transition, with only one or two points along the slope. Due to the micromolar concentrations required for ITC, this technique is generally limited to detecting K_d_ values greater than ∼10^−8^ M, so the values obtained for these interactions must be treated carefully, as they are at this limit.^51, 52^ Whilst the interaction stoichiometries are well-defined, the sharp transition of the binding curves suggests that the true K_d_ values are at or below this detection limit, and therefore should be confirmed with another technique. Accordingly, while thermodynamic parameters are included in the supplementary information (Table S1), they should be interpreted with caution due to these limitations.

To validate the ITC results, microscale thermophoresis (MST) was used to independently assess the binding affinity of these compounds to the 3WJ (Figs. 5A, S20). MST is a highly sensitive technique suited to measuring high affinity interactions. Firstly, the Pt helicate was serially diluted from 5 µM to 153 pM and mixed with a fluorescently labelled, pre-folded 3WJ (Cy5-3WJ-T_3_) (20 nM), in buffer (10 mM Na cacodylate, 100 mM NaCl, 0.1% Tween). The resultant binding curve gave K_d_ = 1.22 ± 0.22 × 10^−8^ M (25 °C). This is consistent with the ITC result. A serial dilution of the Ni cylinder in this range did not provide a suitable binding curve, so it was instead serially diluted from 20 nM to 305 fM and mixed with 3WJ (1 nM; reduced as the DNA concentration should be close to or below the K_d_), revealing K_d_ = 5.15 ± 0.67 × 10^−10^ M — a ∼24-fold higher affinity than the Pt helicate. This confirms that the perfect fit of the Ni cylinder in the 3WJ cavity is better than the induced fit of the Pt helicate, though both are very high affinity interactions.

**Figure 5.**
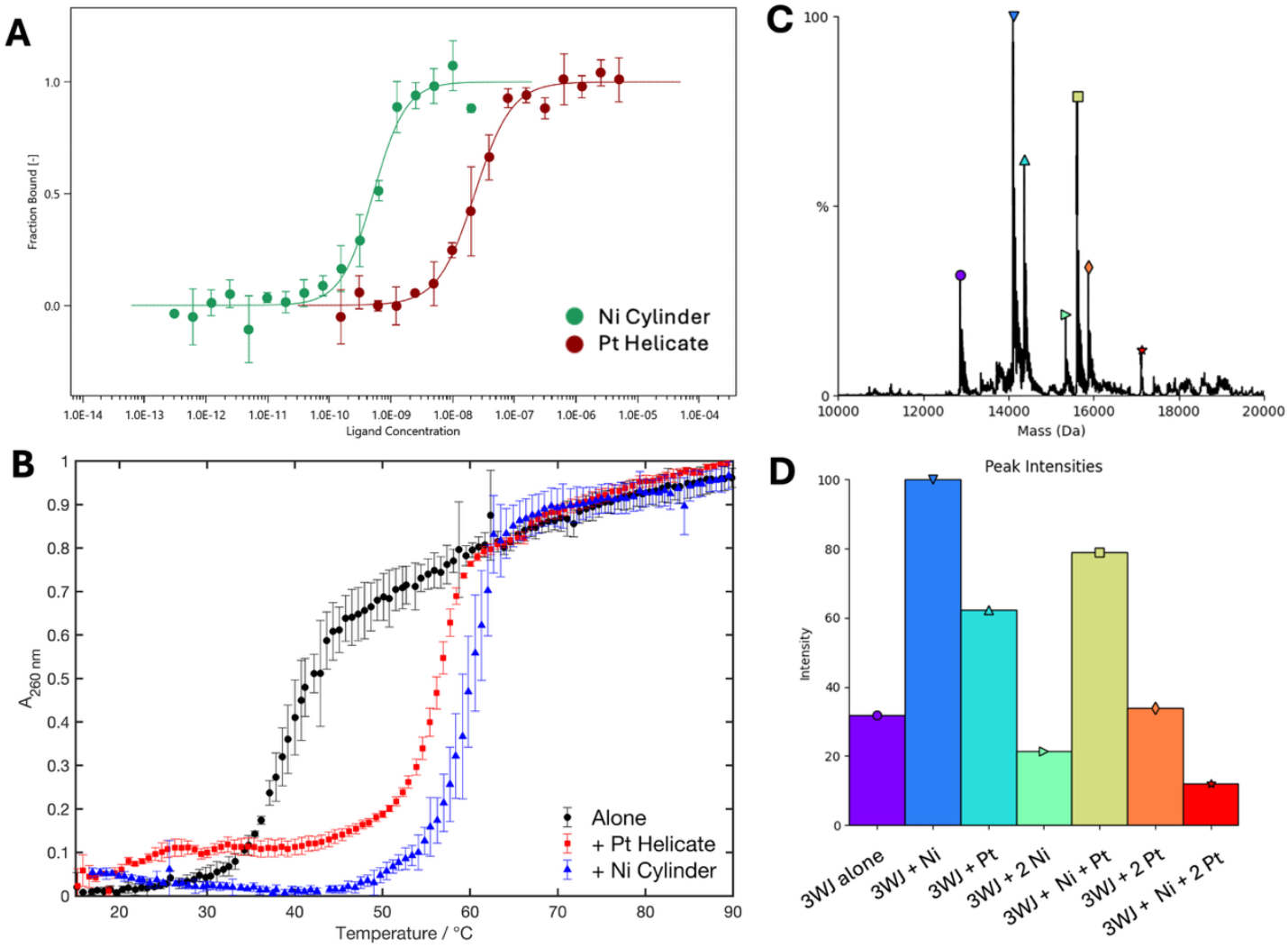
A) Binding curves generated from MST experiments of Cy5–3WJ-T_3_ with Pt helicate (red) and with Ni cylinder (green) (10 mM Na cacodylate, 100 mM NaCl, 0.1% Tween-20). B) UV melting curves of the 3WJ alone (green), and with 1 equiv. Pt helicate (red) or Ni cylinder (purple). Each data point is the average of three independent repeats with associated standard deviation. C) Deconvoluted mass spectrum of the 3WJ exposed to 1 equivalent of Ni cylinder and 1 equivalent of Pt helicate. D) Bar plot of the intensities of the major peaks observed in the MS.

The use of MST here successfully circumvents the detection limit of ITC and captures the picomolar affinity of the cylinder, though is unable to deliver all thermodynamic parameters. As illustration of the strength of 3WJ binding, we highlight that for duplex DNA binding such as the intercalators ethidium, daunorubicin and doxorubicin (clinical drug) and the minor groove binder Hoechst, typical affinities are (at best) in the micromolar range.^53–55^

The strong binding of the compounds is also reflected in UV melting curves (Fig. 5B). The 3WJ alone gave a melting temperature (T_m_) = 39.2 ± 0.9 °C. Upon addition of 1 equivalent Pt helicate, a change (ΔT_m_) of 17.3 ± 1.5 °C was observed. The Ni cylinder led to a slightly larger increase (ΔT_m_ = 20.8 ± 1.6 °C). Whilst melting temperatures are not direct analogues for K_d_, these results are consistent with the stronger binding of the Ni cylinder leading to better stabilisation of the 3WJ compared to the Pt helicate.

We also explored whether native mass spectrometry (MS) could be used to study the 3WJ binding of the Pt helicate and Ni cylinder. The challenge in this technique with our system is whether the peak arises from a bound-drug species, or if the complex and DNA simply fly as a cation–anion pair, however the technique has been used to study G-quadruplexes and recently to detect 3WJs.^56–58^ A pre-folded single-stranded 3WJ (3WJ-T_6_) was first exposed to either 1 equiv. Ni cylinder or 1 equiv. Pt helicate (10 mM NH_4_OAc, 10% MeOH) to confirm that a DNA–complex species could be detected (Figs. S21A, S21B). As DNA can adopt a large number of charge states, deconvolution of the spectrum was used. In both cases, the deconvoluted spectra revealed peaks corresponding to the free DNA and the 1:1 DNA–complex species (Fig. S22). In the case of the Pt helicate, a 1:2 species was also detected. Given the solution 1:1 binding observed in ITC, one of those Pt helicates is likely bound as a simple ion-pair. The 3WJ-T_6_ was then incubated with equal amounts of Ni cylinder and Pt helicate (1 equiv. each) for 1 hour before injecting into ESI-MS (Fig. S21C). After deconvolution, many peaks were present comprising 3WJ + multiple compounds (Fig. 5C). The most intense peak corresponded to the 3WJ + Ni cylinder (Fig. 5D), which may reflect the cylinder’s higher 3WJ affinity, but it seems this is not a straightforward technique to assess relative affinity of these compounds, as the harsh conditions of the ESI-MS injection are not representative of solution behaviour.

### Strength of 4WJ Binding

Despite a strong affinity and clear preference for 3WJs, the PAGE results show that the compound can still bind to 4WJs. UV melting confirmed binding to and stabilisation of the 4WJ (Fig. 6A). Alone in buffer (10 mM Na cacodylate, 100 mM NaCl), the 4WJ melts at T_m_ = 45.7 ± 0.5 °C. On addition of 1 equivalent Pt helicate, T_m_ = 51.7 ± 0.6 °C (ΔT_m_ = 6.0 ± 1.1 °C). In comparison, the Ni cylinder gave T_m_ = 54.5 ± 0.5 °C (ΔT_m_ = 8.8 ± 1.0 °C).

**Figure 6.**
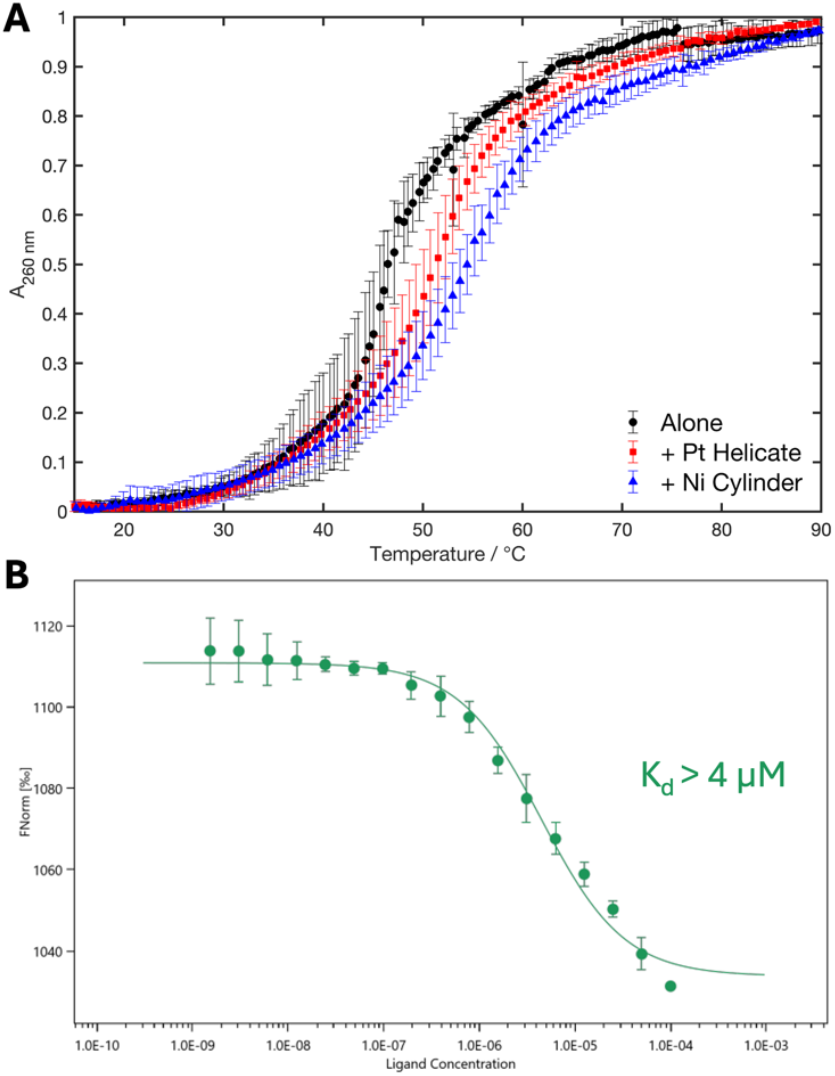
A) Exemplar UV melting curves of the 4WJ alone (green), and with 1 equiv. Pt helicate (red) or Ni cylinder (purple) (10 mM Na cacodylate, 100 mM NaCl). B) Binding curve generated from MST experiments of FAM–4WJ with Pt helicate (10 mM Na cacodylate, 100 mM NaCl).

MST was used again to assess the binding strength of the Pt helicate to 4WJs (Figs. 6B, S23) — we recently reported an equivalent experiment with the Pt-BIMA 4WJ binder, observing a K_d_ of 82 nM.^27^ A FAM-labelled 4WJ was mixed with a serial dilution of the Pt helicate from 100 µM to 1.53 nM, and though the resultant curve was not fully saturated at the higher concentrations, the binding constant is observed to be K_d_ > 4 µM. This is at least two orders of magnitude weaker than the 4WJ binding of Pt-BIMA and nearly ∼400-fold weaker than its binding to the 3WJ.

Simulations were conducted with the 4WJ and were started with the helicate directly inside the open cavity (3 simulations per enantiomer, 3 µs each). In 3 simulations, the junction formed all four branchpoint base pairs around the helicate, though notably the helicate appears to be too small to maintain *π* contacts on all 4 sides simultaneously, causing it to rotate sporadically inside the cavity and unable to adopt one stable binding mode (Fig. S24). Additionally, the cavity here is rhombus shaped (Fig. 7A), as opposed to the square shape exhibited when bound by the Pt-BIMA metallo-cage (Fig. 1B). One simulation showed base pair fraying at one of the branchpoints, leaving a reduced cavity size and a binding pocket consisting of 3 base pairs and one lone base (Fig. 7B), whilst the other two simulations showed distortion of the 4WJ near the cavity as multiple branchpoint base pairs were broken (Fig. 7C), with the helicate unable to find a binding pocket that can stabilise any one DNA conformation. Rarely, chiral inversion of the helicate was observed, but this did not seem to be associated with a stable binding mode, as it was in the case of the 3WJ.

**Figure 7.**
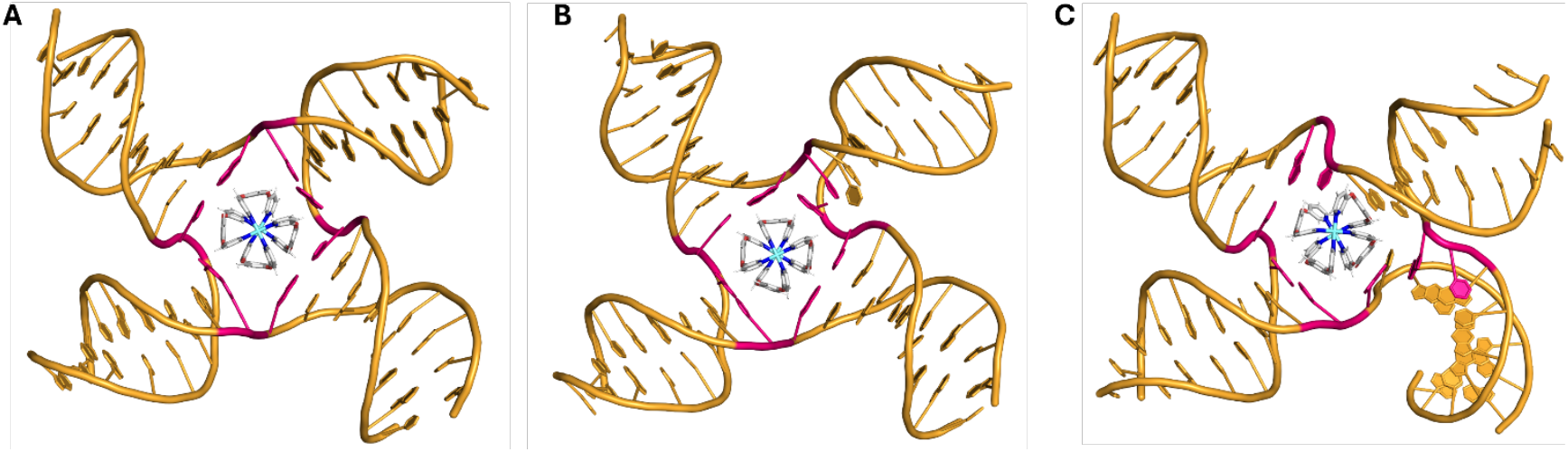
3 MD snapshots of the Pt helicate inside a DNA 4WJ exhibiting A) a rhombus shaped cavity with all 4 base pairs intact; B) a reduced cavity where one base pair has frayed, providing flexibility for the cavity to compress around the helicate; C) a distorted cavity, where fraying of multiple base pairs has caused large dynamic variability in the DNA conformation and partial unwinding of a duplex arm. In all simulations with the 4WJ, the Pt helicate was unable to adopt a stable position inside the cavity and is thus seen to rotate sporadically or cause deformations of the 4WJ structure. Branchpoint base pairs are shown in pink.

These simulations provide some insight into the lower affinity of this compound for the 4WJ, despite its compatible symmetry: it is clear that there is a suboptimal size match between the helicate and the 4WJ cavity, as evidenced by the various possibilities exhibited by simulations beginning from the same starting coordinates. The helicate often struggles to maintain a stable binding mode, and the 4WJ is either heavily distorted or forced to compress its cavity to accommodate for the smaller size of the helicate. Nevertheless, and importantly, the helicate always remained inside the cavity and was able to find metastable states that persisted for hundreds of nanoseconds.

A contributing factor to its very dynamic binding may be that the single, central phenyl moiety on the helicate surface is inappropriately positioned for optimal stacking with the base pairs – that is, it finds itself situated underneath the base pair hydrogen bonds rather than the bases themselves (unlike in the Pt-BIMA metallo-cage, which presents anthracenes that span across the length of the base pair) and is thus not oriented well for *π*-stacking. Binding-induced chiral inversion was not observed in any simulations with the 4WJ, regardless of the starting enantiomer, and this is consistent with a much weaker induced CD signal (Fig. S25).

These simulations also provide some extra insight into why the 3WJ is preferred: the binding to the 3WJ is bolstered by the fact that the hydrophobic 3WJ cavity is innately smaller (so less deformation of the overall DNA structure is required), and that the base pair fraying makes a nucleobase available and well-positioned to stack with a pyridyl coordinating unit, as well as positioning other stabilising interactions in appropriate places.

### Weak Interactions with Double-Stranded DNA

Though DNA junctions are the target of these junction binding metal complexes, we sought also to explore the interaction with dsDNA given that the majority of cellular DNA is found as simple double-stranded B-DNA. While these complexes strongly prefer 3WJs, we have previously shown that the 3WJ-binding triple helicate cylinders induce coiling of DNA with natural biopolymer genomic DNAs such as calf thymus DNA (ctDNA) and linearised plasmids.^59, 60^ We started with simple displacement assays where ctDNA was loaded with either ethidium bromide (EtBr) (an intercalator) or Hoechst 33258 (a minor groove binder) and Pt helicate added. As the concentration of Pt helicate was increased, a gradual decrease in luminescence for both EtBr (IC_50_ > 50 µM, Fig. 8A) and Hoechst (IC_50_ = 8.7 µM, Fig. 8B) was observed. Analogous experiments with the Ni cylinder (Figs. 8C–8D) yielded lower IC_50_ values of 3.4 µM and 3.8 µM (consistent with previous findings^59-61^), indicating that the Ni cylinder binds dsDNA more strongly than the Pt helicate. No changes in the MST traces were observed over a titration of the Pt helicate with a dsDNA oligomer (DS-21) in the range 50 µM to 1.53 nM, consistent with a much weaker binding of the compound to the dsDNA structure compared to the junctions, suggesting a K_d_ at best in the high micromolar range (Fig. S26).

**Figure 8.**
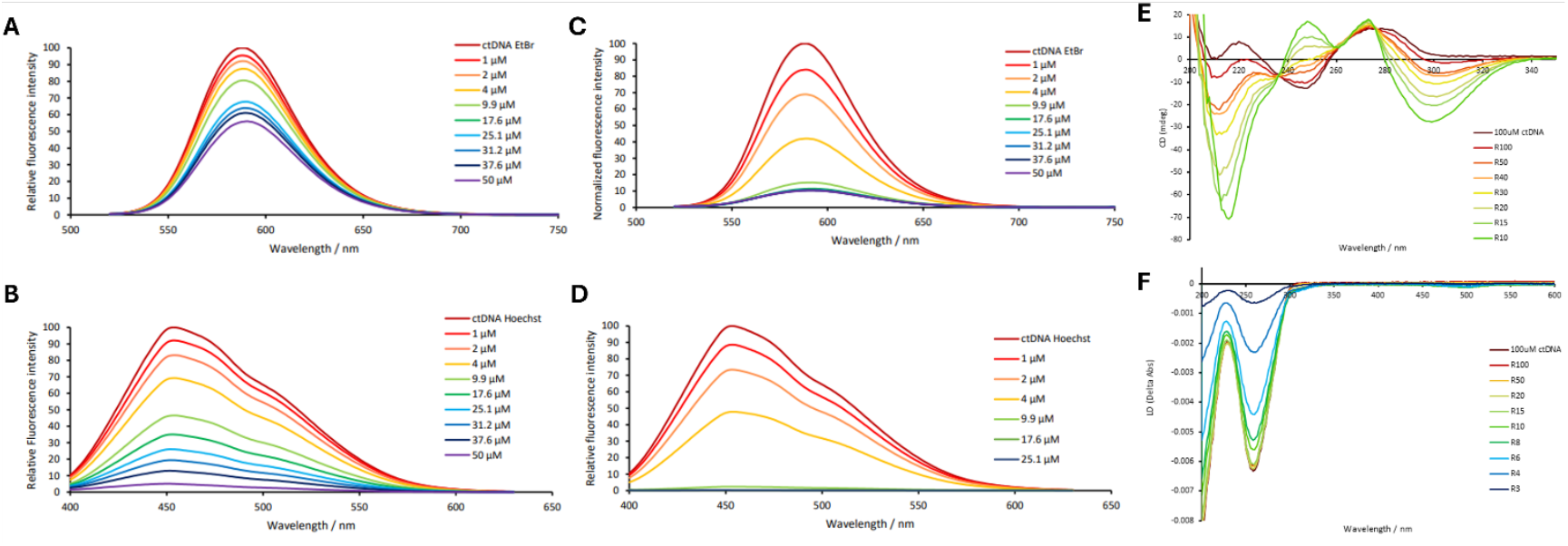
A) EtBr fluorescence displacement titration with Pt helicate; B) Hoechst 33258 fluorescence displacement titration with Pt helicate; C)EtBr fluorescence displacement titration with Ni cylinder; D) Hoechst 33258 fluorescence displacement titration with Ni cylinder. All 50 µM ctDNA, 25 μM EtBr or Hoechst 33258 in 1 mM Tris-HCl, pH 7.4, 20 mM NaCl. E) Circular dichroism and F) linear dichroism titrations of Pt helicate with ctDNA (100 μM base pairs; 1 mM Tris-HCl, 20 mM NaCl, pH 7.4). The R number is the ratio of DNA base pairs:helicate.

CD studies in which the Pt helicate is titrated into ctDNA (Fig. 8E) reveal immediate and dramatic changes in the characteristic B-DNA CD spectrum. There is the formation of a new negative peak at 300 nm which tails out to 340nm, positive peaks at 270 nm and 250 nm as well as a large negative peak at 220 nm. Subtracting the initial DNA CD spectrum reveals the induced CD signals (Fig. S25B), and overlaying these peaks with the induced peaks observed with the 3WJ reveals that the induced CD peaks are very similar in both cases, indicating a preferential binding of the same enantiomer. The CD spectrum of DS-21 with 1 equivalent Pt helicate showed only a weak induced CD band at 290–340 nm suggesting a much weaker interaction (Fig. S25); genomic DNA potentially contains (or can form) non-canonical DNA structures.

To probe whether the Pt helicate bends or coils genomic DNA as the Ni cylinder, linear dichroism was employed. Flow linear dichroism (LD) uses a Couette flow cell to orient the DNA polymer in the direction of flow and then measures the difference in absorbance between linearly polarized light parallel and perpendicular to the direction of flow.^62, 63^ ctDNA shows a characteristic negative band at 260 nm, which gradually reduces in magnitude on addition of increasing concentrations of Pt helicate (Fig. 8F), indicating a loss of orientation consistent with DNA bending or coiling induced by the compound. This phenomenon is also observed when the cylinder is introduced to ctDNA.^59^ The compound alone is too small to be oriented by viscous drag and yield an LD signal, however if binding occurs in a specific orientation, induced LD bands can appear. In this case, a very small induced band appears at around 500 nm, as well as a small shoulder at around 310 nm, indicating the compound is orientated upon binding.

MD simulations of a 25mer B-DNA in the presence of multiple Pt helicates captured the helicate binding transiently along the major and minor grooves of the helix (Fig. S27). Minor groove binding modes persisted longer than binding in the major groove, and were accompanied by a widening of the groove. The helicate was also observed to slide dynamically along the minor groove. Overall, the residence time of all binding events was low, and dissociation occurred readily implying that these interactions are weak and likely arising primarily due to electrostatic attraction between the cationic helicate and the anionic DNA, as opposed to a specific interaction governed by supramolecular geometry. Additionally, helicates were observed to cap the termini of the duplex where the helicate caused fraying of the terminal base pair, allowing stacking to occur with the underlying base pair in addition to the newly opened base pair. Some inversion of chirality was observed in the helicate, however it did not appear to be related to binding and seemed to occur randomly in solution.

## CONCLUSION

This work affords new insight into the key factors that are important for effective junction binding. The new platinum(II) quadruple-stranded cylinder has four-fold symmetry, and so might be expected to be complementary to the 4WJ cavity. Yet MST results show that this helicate has a very strong, (∼10 nM) affinity for 3WJs and a >25 fold weaker affinity for 4WJs. The 3WJ preference is further confirmed by gel competition assays and the tendency of the helicate to induce p*seudo*-3WJs in 4WJ gels. MD simulations suggest that the helicate is slightly too small for an optimal fit in the 4WJ junction cavity, and in fact finds a more suitable binding pocket upon an induced fit in the 3WJ. This suggests that the size-fit is more important than precise shape-fit.

Intriguingly the fit of this platinum quadruple-stranded helicate in the ‘perfect’ (fully base-paired) 3WJ is poor and the helicate causes the 3WJ cavity to rearrange to better accommodate it, opening up a base pair and pushing a base out of the 3WJ cavity to accommodate it (in simulations and consistent with gel results). While this reinforces the dynamic and flexible nature of DNA junction cavities, alongside this the simulations also suggest that the flexible nature of this helicate is vital to achieving the induced fit – with not only the DNA host but the helicate guest also modifying its shape, with inversion of helicity to get the best fit. This indicates that shape-fit is important too.

This contrasts with the triple-helical cylinders that have been characterised crystallographically in the centre of the 3WJ, and the simulations of the Pt-BIMA quadruple-helical complex in the heart of the 4WJ. In those cases, one of the enantiomers is a perfect fit for the junction (lock and key binding) but both enantiomers bind well. Those structures are less flexible than this Pt helicate and show no evidence of enantiomer inversion or other structural changes on binding to the DNA cavity. Consistent with the ‘lock and key’ binding and lack of structure pertubation, the MST results show the cylinder binds 3WJ more than an order of magnitude more strongly than the new quadruple-helicate.

Granzham, Monchaud and colleagues have explored the 3WJ-binding of a variety of tetracationic organic azacryptands containing three aryls in the cryptand arms. The lack of bound metals means that these cryptands are also flexible structures.^64^ Chéron’s MD simulations in that work, suggest that bases at the cavity can unpair and a base insert into the cryptand inside the cavity, giving rise to a change in both cavity and guest structure. That insertion process is dynamic and reversible but may represent another (different) form of induced fit 3WJ-binding (with a base flipping into rather than out of the cavity). Equilibrium dialysis on a related azacryptand indicated a K_d_ for 3WJ in the micromolar range^65^; the newer azacryptands are predicted to bind a little more strongly.^66^

Vázquez, Vázquez and colleagues have explored the 3WJ-binding of tetra- and hexa-cationic peptide triple-helicates.^67^ Those metallo-helicates contain bipyridine ligands linked by a CONHCH_2_CH_2_CONH spacer. They lack the shape-fit and outward-facing π-surfaces of the triple-helical cylinders but will have a similar size. They have reported K_d_ for 3WJ in the range 200–500nM by fluorescence quenching^67, 68^ and, using MST, Vázquez and Kellett have reported K_d_ of 30nM.^69^ The ability of these peptide helicates to recognise and bind the 3WJ despite lacking an optimal shape fit parallels the observations made here for the quadruple helicate. Thus, when the shape complementarity is suboptimal, the size match becomes the dominant factor for binding. In our case, this is aided by induced-fit.

Our previous observations have often stressed the importance of both the size and shape of supramolecular DNA junction binders for binding inside the junction cavity.^18, 26, 27, 47^ The work herein, brings nuance to that analysis. Size-fit seems more important than shape-fit in determining the preferred junction of the binder. But best affinity and selectivity is achieved when both size-fit and shape-fit come together. Nevertheless, whilst a complementary fit is clearly the ideal scenario — reflected in the picomolar affinity of the triple helicate cylinder for the 3WJ — this work demonstrates that those compounds without a perfect match should not be immediately discounted as a potential binder. The affinity of this four-stranded compound for a three-stranded junction shows that the binding of supramolecular architectures to DNA junctions can be through a ‘hand and glove’ scenario, and not only the ‘lock and key’ scenario seen with the cylinders. This opens up new possibilities for designs that can be exploited to fine-tune future DNA and RNA junction-binding compounds.

## ASSOCIATED CONTENT

### Supporting Information

Experimental details, Supplementary Figures S1–S27 and Supplementary Table S1 (PDF). Movies of representative MD simulations illustrated in Figures 3A, S15–S16 (MPEG). The original data supporting the findings in this publication has been placed at UBIRA: https://doi.org/10.25500/edata.bham.00001327.

The authors have cited additional references in the supporting information.^70–82^

## ACKNOWLEDGMENTS

This work was funded by the EPSRC Physical Sciences for Health Centre (EP/L016346/1), the BBSRC Midlands Integrative Biosciences Training Partnership (BB/T00746X/1) and the University of Birmingham, and undertaken in the context of the International Research Network on Nucleic Acid Junctions funded by the CNRS-Chimie (France). The authors acknowledge use of the University of Birmingham Analytical Chemistry and Nano Materials facilities, and the MST facilities provided by the Bio-Analytical Shared Resource Laboratories within the School of Life Sciences, University of Warwick. All simulations were performed using the University of Birmingham BlueBEAR and CaStLeS HPC facilities. The authors thank Prof Richard Napier (U. Warwick) for technical support with MST and Dr Subhendu Karmakar (U. Birmingham) for useful discussions and advice.

