## Supplementary Information for "A Flexible Quadruple-Stranded Helicate Demonstrates a Strong Binding Preference for DNA Three-Way Junctions by Induced-Fit"

In addition to the Supplementary Information herein, all original data and videos for the simulations are also made available online at: https://doi.org/10.25500/edata.bham.00001327.

### TABLE OF CONTENTS


Contributions 2
Experimental Methods 2
- Synthesis of 1,4-Bis(3-pyridyloxy)benzene (L1) 2
- Synthesis of $[Pd_2(L1)_4](BF_4)_4$ 3
- Synthesis of $Pt(DMSO)_2Cl_2$ 3
- Synthesis of $[Pt_2(L1)_4](NO_3)_4$ 3
- **Figure S1.** $^1$H NMR spectrum for $[Pd_2(L1)_4](BF_4)_4$ 4
- **Figure S2.** COSY spectrum for $[Pd_2(L1)_4](BF_4)_4$ 4
- **Figure S3.** ESI-MS spectrum for $[Pd_2(L1)_4](BF_4)_4$ 5
- **Figure S4.** $^1$H NMR spectrum for $[Pt_2(L1)_4](NO_3)_4$ 5
- **Figure S5.** COSY NMR spectrum for $[Pt_2(L1)_4](NO_3)_4$ 6
- **Figure S6.** $^{13}$C NMR spectrum for $[Pt_2(L1)_4](NO_3)_4$ 6
- **Figure S7.** HSQC spectrum for $[Pt_2(L1)_4](NO_3)_4$ 7
- **Figure S8.** nESI-MS spectrum for $[Pt_2(L1)_4](NO_3)_4$ 7
- DNA Sequences 8
- Polayacrylamide Gel Electrophoresis (PAGE) 8
- PAGE Competition Experiments 8
- UV-Visible Spectroscopy 9
- UV Melting 9
- Isothermal Titration Calorimetry (ITC) 9
- Microscale Thermophoresis (MST) 9
- DNA Mass Spectrometry (ESI-MS) 10
- Fluorescence Displacement Titrations 10

Molecular Dynamics Simulations 11
- Parameterisation of the Pt helicate 11
- Parameterisation of DNA 11
- Simulations 11

Supplementary Data 12
- **Figure S9.** UV-VIS spectra of the Pd and Pt helicates 12
- **Figure S10.** UV-VIS spectra of the Pd helicate over time 12
- **Figure S11.** UV-VIS spectra of the Pt helicate over time 12
- **Figure S12.** Representative PAGE competition gels 13
- **Figure S13.** PAGE competition curves for Pt-BIMA 14
- **Figure S14.** Representative Pt-BIMA PAGE competition gels 14
- **Figure S15.** MD RMSD plots for 3WJ + *P* Pt helicate 15
- **Figure S16.** MD RMSD plots for 3WJ + *M* Pt helicate 16
- **Figure S17.** MD starting positions and snapshots for 3WJ simulations 17
- **Figure S18.** ITC data for the Pt helicate with 3WJ 18
- **Figure S19.** ITC data for the Ni cylinder with 3WJ 18
- **Figure S20.** MST traces for the 3WJ + Ni cylinder and Pt helicate 19
- **Figure S21.** Raw mass spectra for the 3WJ MS studies 19
- **Figure S22.** Mass spectra of the pre-folded 3WJ + Ni cylinder or Pt helicate 20
- **Figure S23.** MST traces for 4WJ + Pt helicate 20
- **Figure S24.** MD RMSD plots for 4WJ + Pt helicate 21
- **Figure S25.** Overlay of CD spectra 22

**Figure S26.** MST traces for dsDNA + Pt helicate 22
**Figure S27.** MD snapshots of dsDNA + Pt helicate 23
**Table S1.** Thermodynamic parameters obtained by MST and ITC 23
References 24

### CONTRIBUTIONS

MJH conceived and supervised the project. HDW synthesised and characterised compounds and undertook gels, UV, CD and LD spectroscopy, ESI-MS experiments and ITC. SJD undertook DFT calculations and MD simulations, MST experiments, gels, UV melting, and CD. SB synthesised compounds used for some of the studies and performed CD. HDW, SJD and MJH analysed the data and wrote the manuscript which all authors discussed and commented on.

### EXPERIMENTAL METHODS

All solvents, NMR solvents, chemical reagents, buffer components and DNA oligos were purchased from Fischer Scientific, VWR chemicals or Sigma Aldrich and used without further purification. Nickel cylinder $[Ni_2L_3]Cl_4$ was prepared as described previously.[1] Pt-BIMA $[Pt_2(BIMA)_4](NO_3)_4$ was synthesised as previously described.[2] Electrospray ionisation (ESI) mass spectrometry characterisation was carried out on a Waters SYNAPT-G2-S in positive ion mode. $^1$H NMR studies were carried out on AVANCE NEO400 and AVIII 400 (400MHz) Bruker spectrometers. Elemental analysis was performed on a CE Instruments EA1110 elemental analyzer. Nanopure™ water was used in all biophysical experiments. Calf-thymus DNA (ctDNA) for B-DNA experiments was dissolved in water at 1 mM (in base pairs, confirmed by UV-VIS spectroscopy using ε = 13100 M $cm^{-1}$ $bp^{-1}$).

#### Synthesis of 1,4-Bis(3-pyridyloxy)benzene (L1)

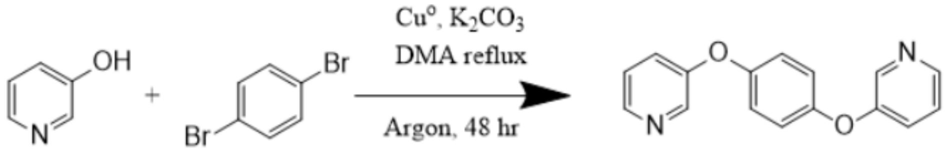


1,4-Bis(3-pyridyloxy)benzene was synthesised as previously reported.[3] 1,4-Dibromobenzene (236 mg, 1 mmol), 3-hydroxypyridine (209 mg, 2.2 mmol), potassium carbonate (607 mg, 4.4 mmol), and copper powder (279 mg, 4.4 mmol) were dissolved in dimethyl acetamide. The reaction mixture was heated under reflux for 48 hr under argon. The solvent was then removed under reduced pressure and ethyl acetate (5 mL) was added. The resulting solution was refluxed for a further 1 hr before the mixture was filtered and washed through with more hot ethyl acetate. The filtrate was concentrated under reduced pressure and purified by silica column chromatography (hexane : ethyl acetate, 50:50 to 100:0). Yield = 172 mg, 65 %.

1H NMR (400 MHz, $CD_3CN$): δ 8.40–8.435 (m, 2H), 8.33 (dd, J = 4.0, 2.0 Hz, 2H), 7.37–7.32 (m, 4H), 7.10 (s, 4H) ppm

1H NMR (400 MHz, *d6*-DMSO): δ 8.42–8.38 (m, 2H), 8.36 (dd, J = 4.1, 1.9 Hz, 2H), 7.47–7.38 (m, 4H), 7.15 (s, 4H) ppm

13C NMR (400 MHz, $CD_3CN$): δ 155.24, 141.71, 153.54, 145.32 ppm

**Synthesis of $[Pd_2(L1)_4](BF_4)_4$**

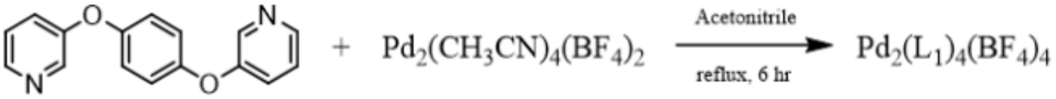


The Pd helicate was synthesized as previously reported.[4] L1 (20.1 mg, 0.076 mmol) was dissolved in acetonitrile (4 mL). $Pd(MeCN)_4(BF_4)_2$ (16.8 mg, 0.038 mmol) was dissolved separately in acetonitrile (1 mL) and added to the ligand solution (leftover remnants were washed in with a further 1 mL acetonitrile). The solution was refluxed for 3 hours before removing the solvent under reduced pressure. The crude product was triturated with diethyl ether (5 mL) before being filtered and washed with more diethyl ether (20 mL). Yield = 24 mg, 78 %.

1H NMR (400 MHz, $CD_3CN$): δ 8.63 (dd, J = 5.6, 1.1 Hz, 8H), 8.57 – 8.53 (m, 8H), 7.86 – 7.79 (m, 8H), 7.65 (dd, J = 8.6 Hz, 5.6 Hz, 8H), 6.74 (s, 16H)

HRMS (ESI-MS) for $[Pd_2(L1)_4(BF_4)]^{3+}$ $C_{64}H_{48}BF_4N_8O_8Pd_2$ = 452.3905 m/z, Found = 452.3930 m/z

**Synthesis of $Pt(DMSO)_2Cl_2$**

This platinum precursor was synthesised as previously reported.[5] $K_2PtCl_4$ (250 mg, 0.6 mmol, 1 equiv.) was dissolved in minimal $H_2O$ (~1.5 mL). To this, DMSO (107 µL, 1.5 mmol, 2.5 equiv.) was then added and the mixture stirred at room temperature for 1 hour. Once an off-white/yellow precipitate began to form, the flask was then placed in the fridge for 4 hours. The resultant off-white precipitate was collected by filtration, washed with cold water, methanol and diethyl ether, and then dried in a vacuum desiccator. Yield = 233 mg, 92%.

$^1$H NMR (400 MHz, *d6*-DMSO) δ 2.54 (s, 6H).

**Synthesis of $[Pt_2(L1)_4](NO_3)_4$**

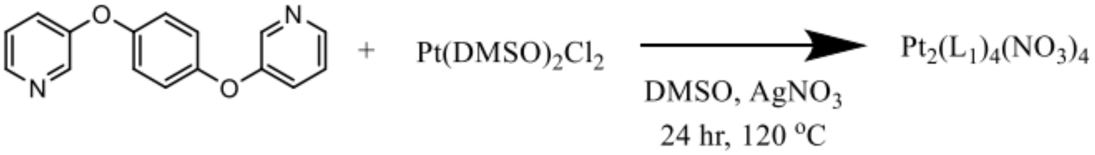


$Pt(DMSO)_2Cl_2$ (40 mg, 0.095 mmol) was dissolved in DMSO (3 mL) under argon. L1 (50.3 mg, 0.19 mmol) was dissolved separately in DMSO (3 mL) before being added to the Pt solution. The flask was washed with a further 3 mL DMSO to ensure maximum recovery. The reaction solution was heated to 120 °C, this temperature was maintained for 24 hr, after which the solution was cooled and filtered through a syringe filter (0.2 uM, PTFE) to remove the AgCl precipitate. The DMSO was then removed by sequential washes and centrifugations with large excesses of diethyl ether (approximately 3 x 30 mL) until the product precipitated. The product was washed once more with diethyl ether before being collected by filtration and dried under high vacuum. Yield = 65 mg, 80 %

1H NMR (400 MHz, *d6*-DMSO): δ 6.77 (s, 16H), 7.85 (dd, J = 8.6, 5.6 Hz, 4H), 8.03 (ddd, J = 8.6, 2.7, 1.1 Hz, 4H), 8.84 (d, J = 2.7 Hz, 4H), 8.89 (dd, J = 5.6, 1.1 Hz, 4H) ppm

13C NMR (400 MHz, *d6*-DMSO): δ 155.07, 151.69, 146.64 119.24, 141.31, 131.58, 128.25 ppm

HRMS (ESI-MS) for $[Pt_2(L1)_4(NO_3)]^{3+}$ $C_{64}H_{48}N_9O_{11}Pt_2$ calculated = 502.7583 m/z, Found = 502.7674 m/z

Elemental analysis: Expected: C 45.34, H 2.85, N9.91 Measured: C 45.41, H 2.85, N9.68

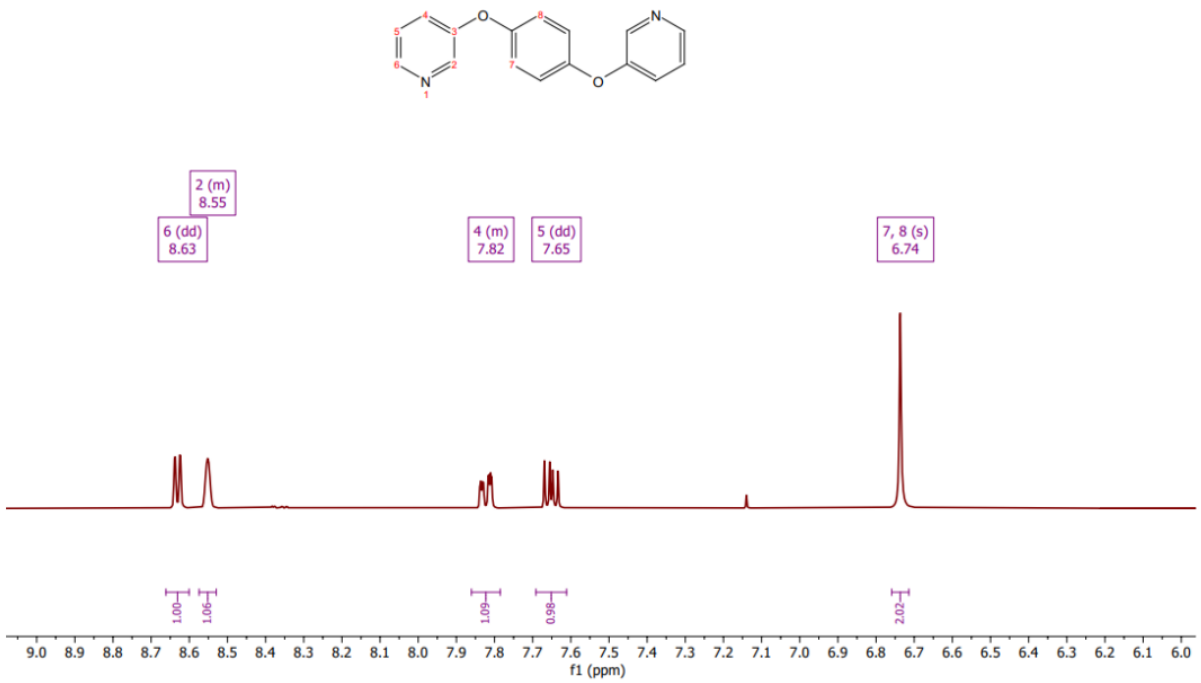


**Figure S1.** [1]H NMR spectrum of $[Pd_2(L1)_4](BF_4)_4$ in $CD_3CN$ (400 MHz).

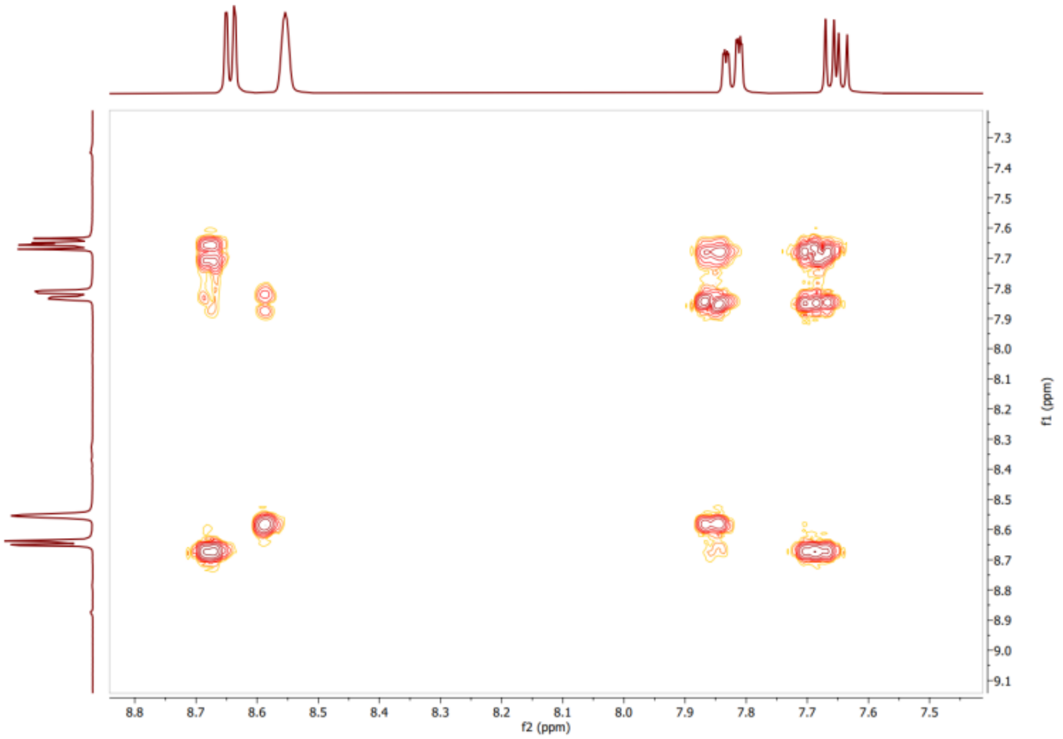


**Figure S2.** COSY NMR spectrum of $[Pd_2(L1)_4](BF_4)_4$ in $CD_3CN$ (400 MHz).

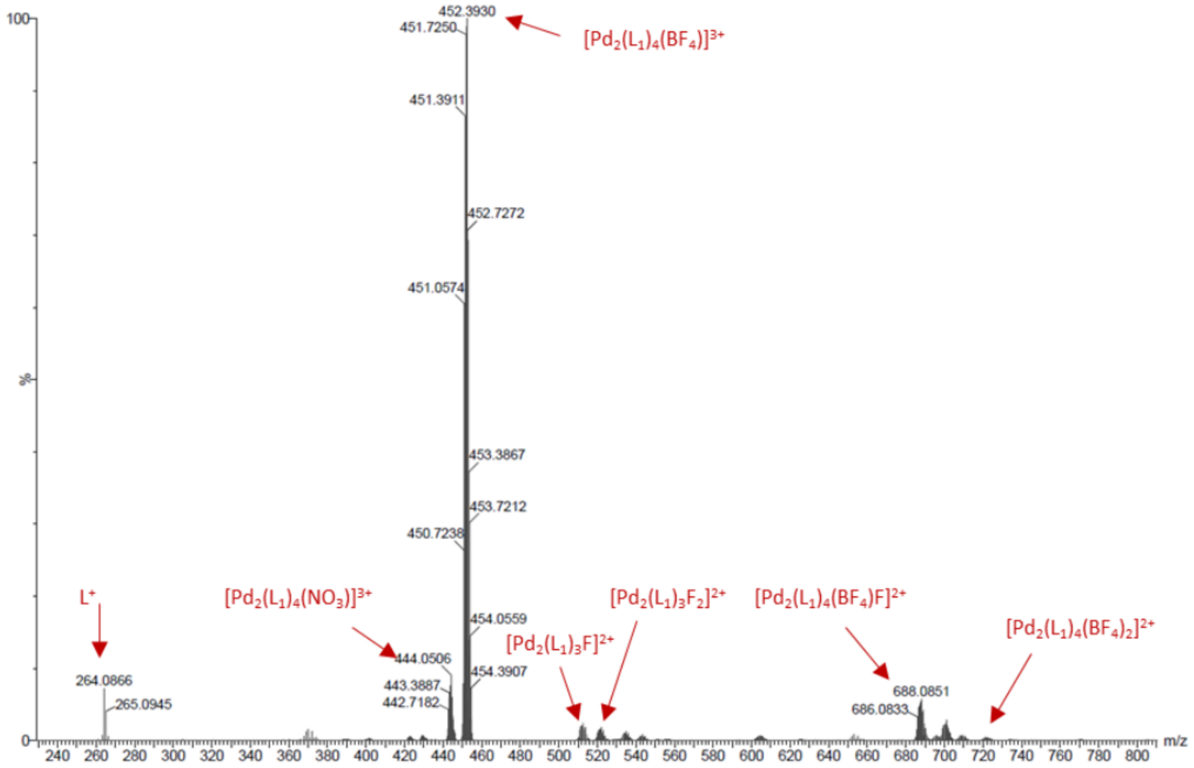


**Figure S3**. ESI mass spectrum of $Pd_2(L1)_4(BF_4)_4$ in ACN. $L1^+$ = 264 m/z, $[Pd_2(L1)_4(BF_4)]^{3+}$ = 452 m/z (Calc. 452.3905), $[Pd_2(L1)_4(NO_3)]^{3+}$ = 443 m/z, $[Pd_2(L1)_4(BF_4)(F)]^{2+}$ = 687 m/z.

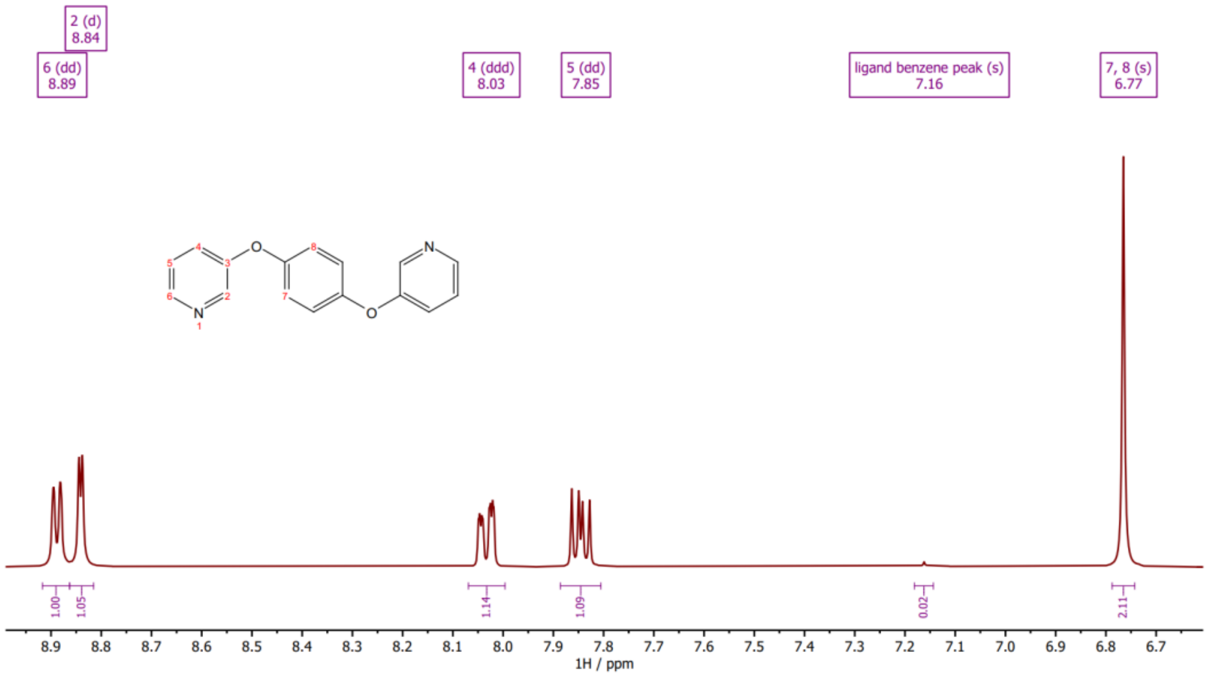


**Figure S4.** $^1$H NMR spectrum of $[Pt_2(L1)_4](NO_3)_4$ in *d6*-DMSO (400 MHz).

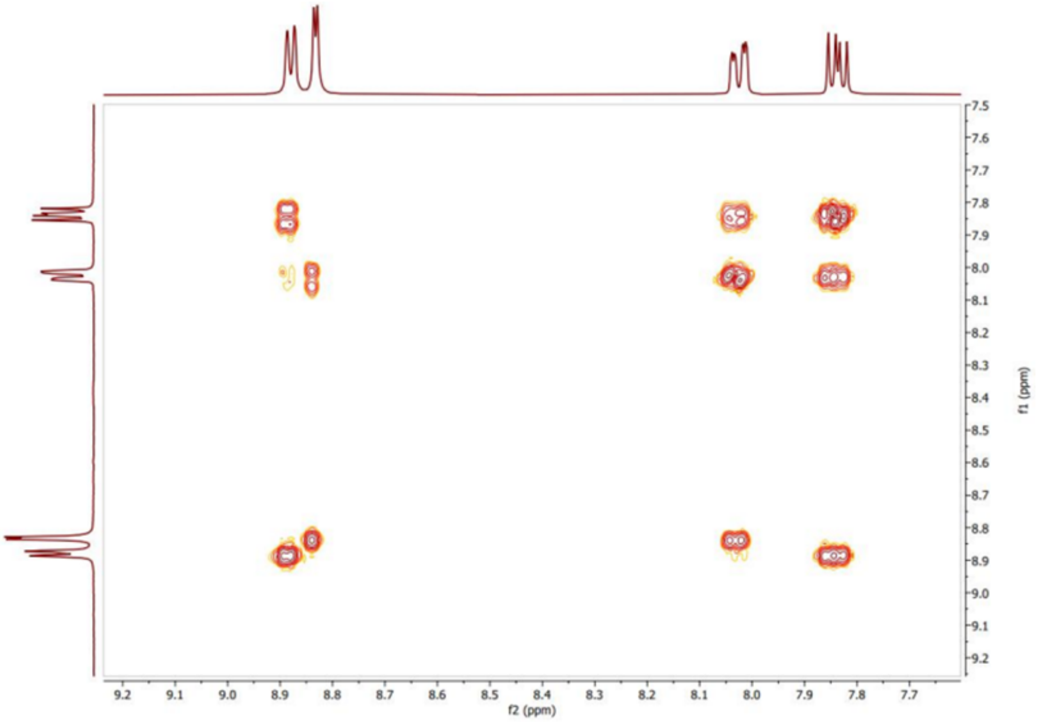


**Figure S5.** COSY NMR spectrum of $[Pt_2(L1)_4](NO_3)_4$ in *d6*-DMSO (400 MHz).

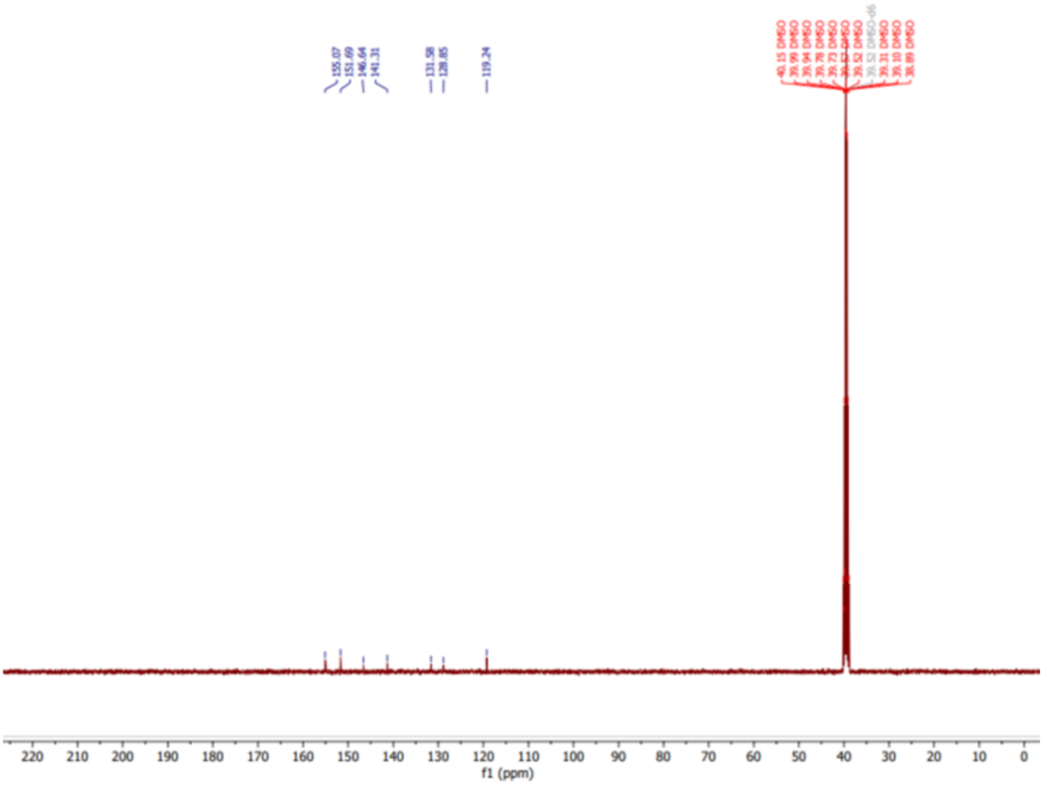


**Figure S6.** $^{13}C$ NMR spectrum of $[Pt_2(L1)_4](NO_3)_4$ in *d6*-DMSO (400 MHz).

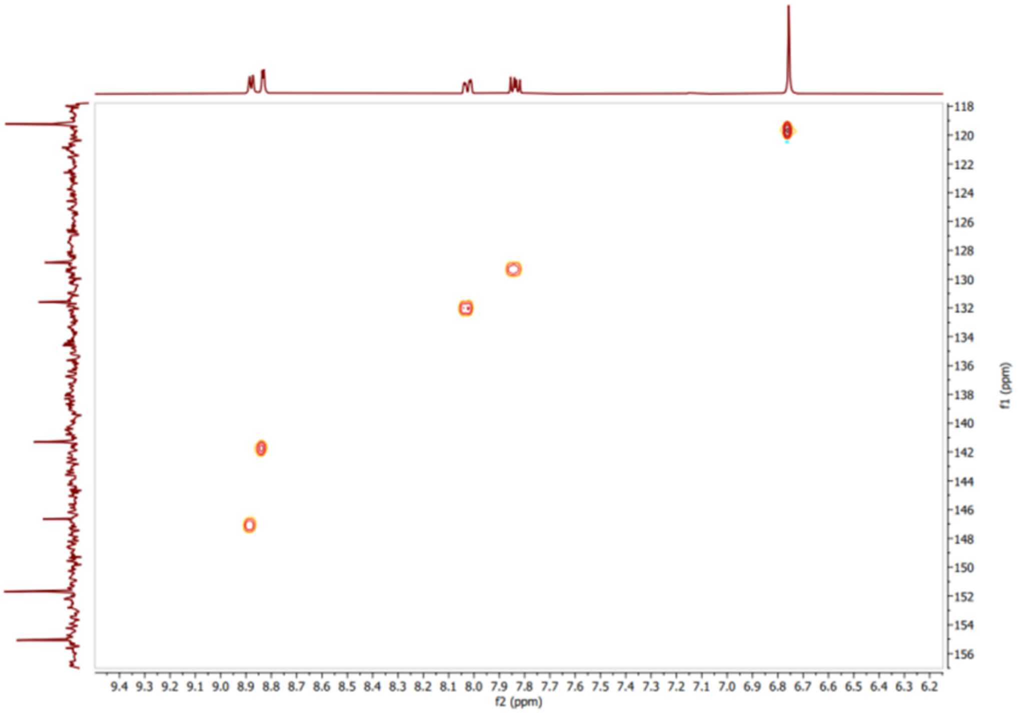


**Figure S7.** HSQC NMR spectrum of $[Pt_2(L1)_4](NO_3)_4$ in *d6*-DMSO (400 MHz).

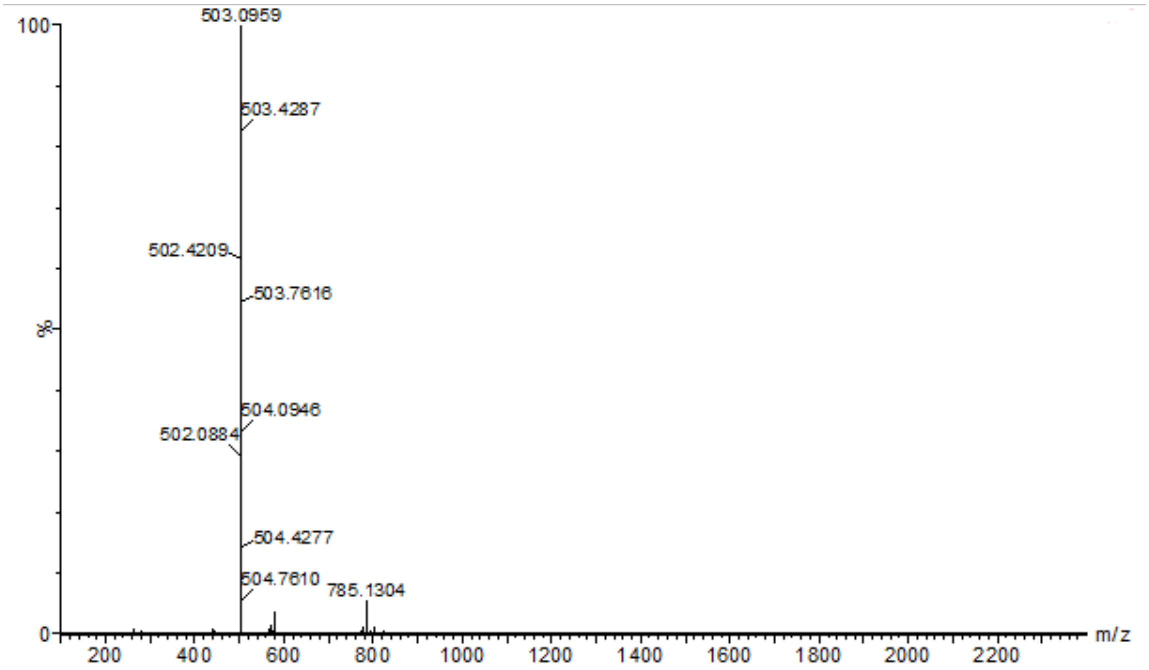


**Figure S8.** ESI mass spectrum of $Pt_2(L1)_4(NO_3)_4$ in MeOH. $[Pt_2(L1)_4(NO_3)]^{3+}$ = 503 m/z (Calc. 503.0928), $[Pt_2(L1)_4(NO_3) + MeOH]^{3+}$ = 785 m/z (Calc. 785.1505).

**DNA Sequences (5′ to 3′)**

DS-21 S1: CCTTCACGCGAACGTAATCCT
DS-21 S2: AGGATTACGTTCGCGTGAAGG

4WJ S1: GCCTAGCATGATACTGCTACCG
4WJ S2: CGGTAGCAGTACCGTTGGTGGC
4WJ S3: GCCACCAACGGCGTCAACTGCC
4WJ S4: GGCAGTTGACGTCATGCTAGGC

Y-fork: 4WJ S2 + 4WJ S3

3WJ S1: CGGAACGGCACTCG
3WJ S2: CGAGTGCAGCGTGG
3WJ S3: CCACGCTCGTTCCG

3WJ-18 S1: GTGGCGAGAGCGACGATC
3WJ-18 S2: GATCGTCGCAGAGTTGAC
3WJ-18 S3: GTCAACTCTTCTCGCCAC

3WJ-$T_3$ (MST): CGGAACGGCACTCGTTTCGAGTGCAGCGTGGTTTCCACGCTCGTTCCG
3WJ-$T_6$ (ITC): ACTCTTCTCGTTTTTTCGAGAGCGACTTTTTTGTCGCAGAGT

**Polyacrylamide Gel Electrophoresis (PAGE)**

20cm x 20cm large 15% PAGE gels were prepared by mixing 25 mL of 37.5:1 acrylamide/bis-acrylamide with 5 mL of 10x Tris-Boric acid buffer (890 mM, pH 8.3) and 20 mL of Milli-Q water. To this 400 µL of a 10% w/v ammonium persulfate solution in water and 40 µL of TEMED were added to initialise polymerisation. This was then immediately poured between 2 glass plates and a 20-well comb inserted at the top; this was then allowed to set for at least 30 minutes before proceeding.

8.3cm x 7.3 cm mini 12% PAGE gels were prepared by mixing 6 mL of 37.5:1 acrylamide/bis-acrylamide with 1.5 mL of 10x Tris-Boric acid (TB) buffer (890 mM, pH 8.3) and 7.5 mL of Milli-Q water. To this 150 µL of a 10% w/v ammonium persulfate solution in water and 15 µL of TEMED were added to initialise polymerisation. This was then immediately poured between 2 glass plates and a 10-well comb inserted at the top; this was then allowed to set for 1 hr before proceeding.

The gel was then attached to the gel jacket and submerged in 1x TB buffer at the top and bottom. The wells were thoroughly flushed before loading of any sample. Samples were made up to 30 µL containing 1 µM of each DNA strand, 1x buffer (89 mM Tris, 89 mM Boric acid, 10 mM NaCl, pH 8.3), and the indicated ratio of complex. DNA, water, and buffer were mixed in solution before addition of the stated ratios of complex. Samples were then centrifuged and incubated at room temperature for 1 hr. 7.5 µL of 50% v/v glycerol was then added to each sample (10% v/v final concentration) and the sample was then centrifuged, mixed, and 12 µL pipetted into the wells on the gel. The gels were run at 140 V for 2.5 hrs (large gels) in 1x TB running buffer. The gel was then removed from the plates and stained using SYBR™ Gold Nucleic Acid Gel Stain (Thermofisher scientific) in 1x TB buffer for at least 30 minutes before imaging on an AlphaImager™ UV transilluminator (Alpha Innotech) with 302 nm excitation.

**PAGE Competition Experiments**

20cm x 20cm 15% PAGE gels (for the Pt helicate competition gels) and 8.3cm x 7.3 cm mini 12% PAGE gels (for the Pt-BIMA competition gels) were prepared as described above. The gel was then attached to the gel jacket and submerged in 1x TB buffer at the top and bottom. The wells were thoroughly flushed before loading of any sample. Samples were made up to 30 µL containing 1 or 2 μM of fluorescently labelled 3WJ (5′ FAM label on strand S1), 1x TBN buffer (89 mM Tris, 89 mM Boric acid, 10 mM NaCl (Pt helicate gels) or 50 mM NaCl (Pt-BIMA gels; to ensure sufficient 3WJ signal), pH 8.3), 1 equivalent compound, and the indicated ratio of competitor DNA (3WJ, Y-fork or DS-21). DNA, water, and buffer were mixed in solution before addition of the stated ratios of complex. Samples were then centrifuged and incubated at room temperature for 1 hr. 7.5 μL of 50% v/v glycerol was then added to each sample (10% v/v final concentration) and the sample subsequently centrifuged, mixed, and 12 μL pipetted into the wells on the gel. The gels were run at 140 V for 1 hour (large gels) or 35 mins (mini gels) in 1x TB running buffer. The gel was then removed from the plates and rinsed in deionised water for 5 minutes before imaging on an AlphaImager™ UV transilluminator (Alpha Innotech) with 302 nm excitation. ImageJ was used to quantify the intensity of the gel bands.[6] The intensities of the 3WJ bands were measured as a fraction of the total lane intensity (ssDNA band intensity + 3WJ band intensity) and normalised to the lane containing no competitor DNA. All competitor ratios were measured as the average of 3 independent samples.

**UV-Visible Spectroscopy**
Samples were dispensed into a 1 cm path length, masked quartz cuvette and absorbance was recorded between 200–800 nm (1 nm bandwidth, 600 nm/min) in a Cary5000 UV-Vis-NIR Spectrophotometer (Agilent Technologies, Inc.) equipped with a multi-cell holder. In all cases, each spectrum was zeroed and a baseline recorded for each condition.

**UV Melting**
Each sample contained 1 μM of each oligo (corresponding to 3WJ-18 or 4WJ), 1 μM of Pt-BIMA, 1% DMSO, 10 mM sodium cacodylate (pH 7.4) and 100 mM NaCl. Control samples were also prepared containing all components except the Pt complex. Samples were dispensed into masked quartz cuvettes with 1 cm path length and the cuvette then stoppered. The measurements were carried out on a Cary5000 UV-Vis-NIR spectrophotometer (bandwidth, 1 nm; average time 1 s; heating rate, 1˚ C $min^{-1}$; measurement interval, 0.5 ˚C) equipped with a multi-cell holder and peltier temperature controller. Data was collected in triplicate for each condition. The data was normalised and the melting temperature ($T_m$) determined as the temperature at the derivative maximum. The final melting temperature was then reported as the average of the three runs with standard deviation error.

**Isothermal Titration Calorimetry (ITC)**
ITC was carried out on a Malvern Microcal PEAQ-ITC instrument. Before using the instrument, a water–water titration calibration was performed. All ITC measurements were carried out in 10 mM sodium cacodylate and 100 mM NaCl (pH 7.4). Stock solutions of the metal complexes in buffer were prepared at a concentration of 150 μM. A DNA stock solution of 3WJT6 (a single stranded oligo that forms 3WJ secondary structure) was annealed by heating at 95° C for 5 minutes then allowed to cool slowly. The DNA solution was then diluted to 10 μM and placed into the cell. The metal complex solution was loaded into the syringe and titrated into the cell. Each titration was repeated 3 times to ensure reproducible results. Controls were also carried out where the metal complex was titrated into buffer and where buffer was titrated into the 3WJT6 solution. It should be noted that the platinum complex required 0.5 % acetonitrile to fully dissolve, thus the amount of acetonitrile was kept consistent throughout the experiment to avoid solvent-mixing effects.

**Microscale Thermophoresis (MST)**
A 5 mM stock solution of the Pt complex in 50% aqueous acetonitrile was diluted to either 200 μM or 10 μM in buffer (10 mM Na cacodylate, 100 mM NaCl, 0.1% Tween). A 1 mM solution of Ni cylinder was diluted to 20 nM in the same buffer. Samples were prepared by making a serial dilution of complex in buffer and mixing with equal volume of either Cy5-labelled 3WJ-T3 (2 nM or 40 nM) or FAM-labelled 4WJ (40 nM). Samples were incubated at room temperature for at least 15 minutes before loading

into Monolith standard capillary tubes and the tubes then placed into a Monolith NT.115 (Nanotemper Technologies). MST experiments were run at room temperature (22 °C) using 60% excitation power (blue laser) and medium MST power. Each experiment was done in triplicate. The data was analysed and plotted using MO.Affinity Analysis software (Nanotemper Technologies).

**DNA Mass Spectrometry**

DNA only samples (10 µL total volume) were prepared at a concentration of 50 µM pre-folded 3WJ in ammonium acetate buffer (10 mM, pH 7.0) in water. Samples were then allowed to equilibrate for 1 hour at room temperature. Samples with the addition of metal complexes were prepared in the same way using 1 mM stock solutions of the metal complexes and reducing the volume of water added. The solutions were then diluted to 10 µM in 10% MeOH immediately before running on the mass spectrometer. MeOH was used in order to achieve a stable spray for the MS measurement. All mass spectrometry was carried out on a Waters SYNAPT G2 with an Advion TriVersa Nanomate® source. Injection method used gas pressure 0.7 psi, Voltage 2.2 kV, source temperature 70 °C, and a cone voltage 80 V. The mass was scanned from 100 – 2400 m/z. Data was processed using UniDec software.[7]

**Fluorescence Displacement Titrations**

Fluorescence displacement experiments were adapted from the procedure reported by Chaoyang Li et al.[8] Fluorescence measurements were taken on a Jasco FP-8500 in a 1 cm quartz cuvette. Ethidium Bromide and DNA stock solution: 100 µM ctDNA (in base pairs), 2 mM Tris-HCl pH 7.4, 40 mM NaCl, and 50 µM Ethidium bromide. Hoechst 33258 and DNA stock solution: 100 µM ctDNA (in base pairs), 2 mM Tris-HCl pH 7.4, 40 mM NaCl, and 10 µM Hoechst 33258. The stock solutions were diluted to a DNA concentration of 50 µM and a volume of 1 mL. The fluoresce of this sample was then recorded before aliquots of the relevant metal complex and an equal volume of the stock solution were added to the working solution. The solution was allowed to equilibrate with gentle stirring for 5 minutes before acquiring the fluorescence measurement. Each measurement was taken three times to ensure the fluorescence intensity of the sample was not changing over time. Parameters for Ethidium bromide fluorescence displacement: $\lambda_{exc}$ = 500 nm, $\lambda_{em}$ = 520 – 750 nm, Data interval = 1 nm, Sensitivity = High, Excitation bandwidth = 2.5 nm. Parameters for Hoechst 33258 fluorescence displacement: $\lambda_{exc}$ = 350 nm, $\lambda_{em}$ = 400–650 nm, Data interval = 1 nm, Sensitivity = Medium, Excitation bandwidth = 2.5 nm.

### MOLECULAR DYNAMICS SIMULATIONS

**Parameterisation of the Pd and Pt helicates.**

Parameters for the coordination bonds were calculated using the MCPB.py pipeline with Gaussian09 at the ωB97XD/DEF2-SVP level of theory to include dispersion, with ECP for Pd and Pt.[9–10] The output coordinate and topology files were converted to GROMACS format using ParmEd (https://github.com/ParmEd/ParmEd). Coordinates for the mirror image enantiomers were generated using the invert chirality function in Avogadro.[11]

**Parameterisation of DNA**

The PDB file for the 25mer B-DNA consisting of 2 strands ($A_{25}$ and $T_{25}$) was generated using NAB (nucleic acid builder) in AmberTools. [12] The 3WJ structure was adapted from PDB 1F44,[13] as described previously.[14] The 4WJ structure was taken from the 1XNS PDB crystal structure,[15] and adapted as described previously.[14] The "true" 4WJ (in which all base pairs are intact from the beginning) was taken from a snapshot of an MD simulation we previously reported containing an organometallic pillarplex in the cavity;[14] the pillarplex was removed leaving the DNA alone. Similarly, the closed 4WJ structure was taken from a snapshot of an MD simulation of the free 4WJ (i.e. with no compound). All DNA was parameterised using the AMBER forcefield parmbsc1.[16]

**Simulations**

MD simulations were carried out as previously described.[2, 14, 17–18] In all simulations, DNA was placed with compound inside a dodecahedral box with edges at least 1.0 nm away from the DNA. All systems were solvated in water using the TIP3P model and neutralised with $Na^+$ ions. Additional $Na^+$ and $Cl^-$ ions were added to reach a NaCl concentration of 50mM. Using GROMACS software[19] initial minimisation was carried to at least 500 kJ/mol/nm or 50000 steps followed by heating and NVT equilibration for 1000 ps using V-rescale modified Berendsen thermostat, coupling the cylinder with the DNA at 310K. All simulations use 2 fs time step and Parrinello–Rahman pressure coupling and PME electrostatics at 1.0nm cut-off. All simulations were run on the BlueBEAR cluster at U. Birmingham using the CaStLeS resources. After the simulations had finished, the trajectories were processed in GROMACS to remove periodic boundary conditions, translations and rotations, and visualised in PyMOL.[20] RMSD plots were calculated using the rms function in GROMACS and plotted using MATLAB.

### SUPPLEMENTARY DATA

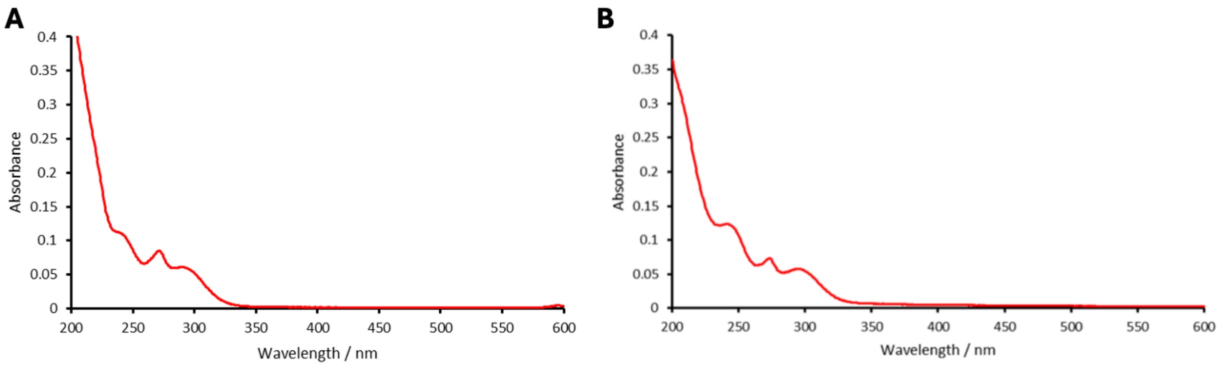


**Figure S9.** Absorbance spectra of $[Pd_2(L1)_4](BF_4)_4$ (A) and $[Pt_2(L1)_4](NO_3)_4$ (B); 2.5 µM in buffer (1 mM Tris-HCl, 20 mM NaCl, 0.05% acetonitrile) recorded at room temperature.

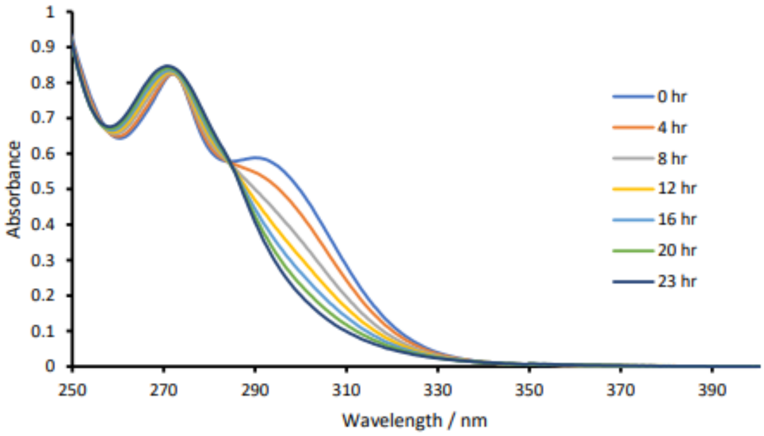


**Figure S10.** UV-Vis absorbance measurements of $Pd_2(L1)_4(BF_4)_4$ 25 µM in buffer (89 mM TB, 10 mM NaCl, pH 8.3) over time with 1% DMSO. The change in the curve over 23 hours reveals degradation of the compound.

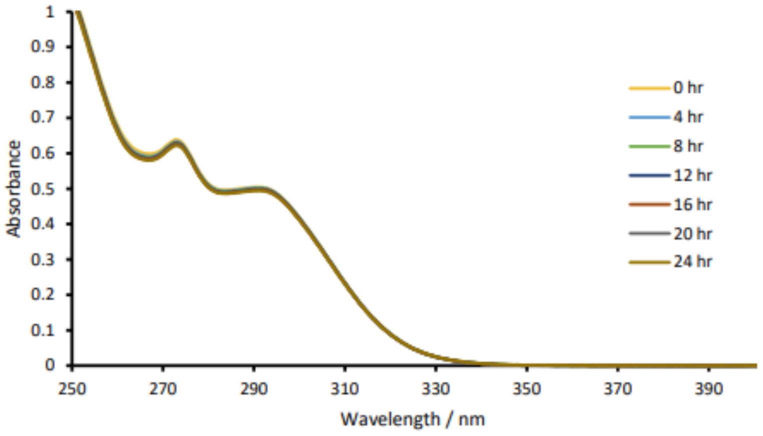


**Figure S11.** UV-Vis absorbance measurements of $Pt_2(L1)_4(NO_3)_4$ 25 µM in Tris-borate buffer 89 mM pH 8.3, NaCl 10 mM over time with 1% DMSO.

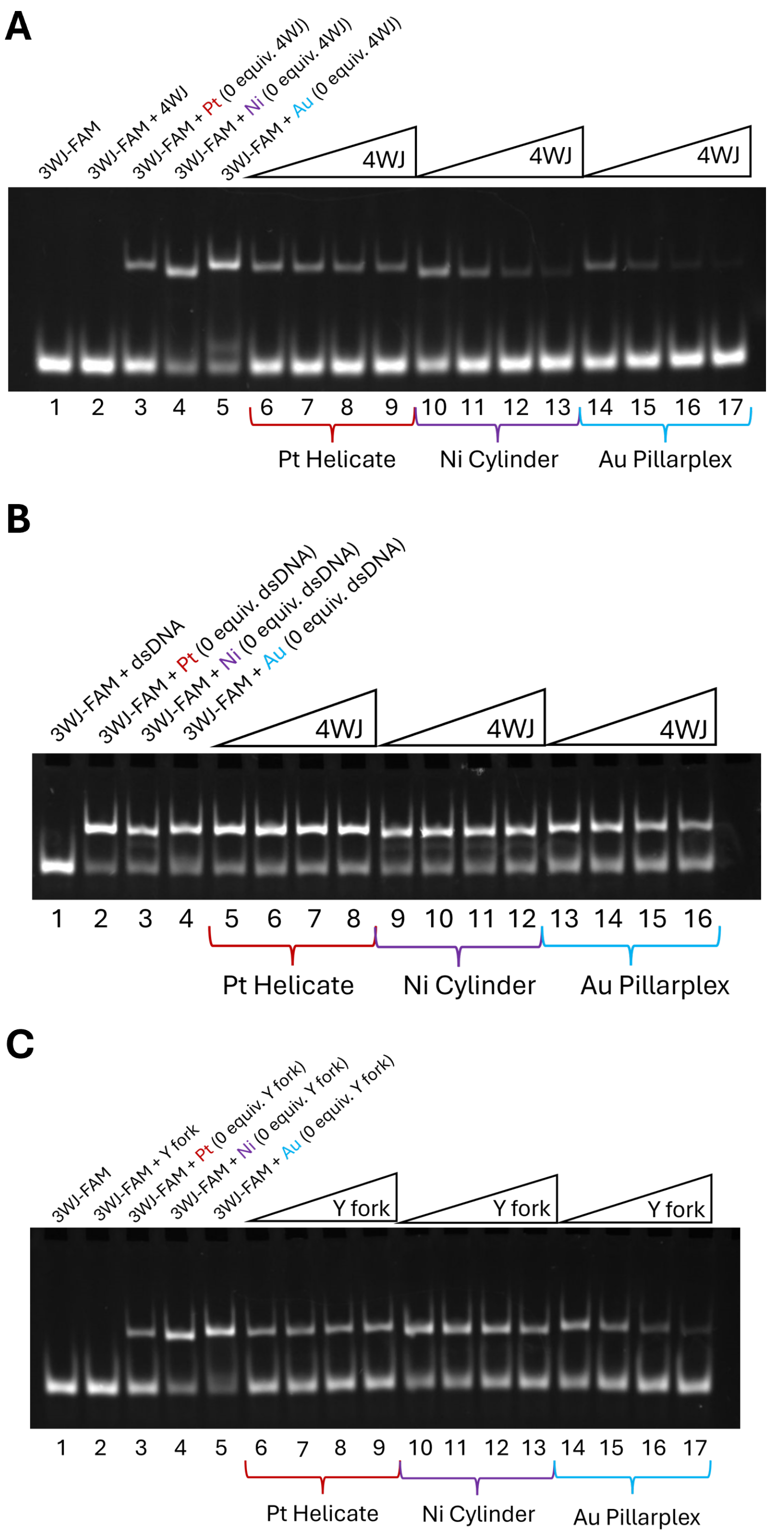


**Figure S12.** Representative gels for the Pt helicate PAGE competition experiments. Each sample contained 1 µM FAM–3WJ and 1 equivalent compound with increasing concentrations of (A) 4WJ, (B) dsDNA or (C) Y fork (0, 0.5, 1, 2 and 4 equivalents). Each gel was repeated 3 times. The intensity of the 3WJ band was measured as a ratio of the total fluorescence in the lane and averaged across the 3 repeats. The plotted curves can be seen in Fig 3B. The first two lanes of (B) have been cropped as they contain samples not discussed in this publication. Lanes pertaining to Ni cylinder or Au pillarplex have been previously published in *J. Am. Chem. Soc.* **2023** *145* (25), 13570–13580.

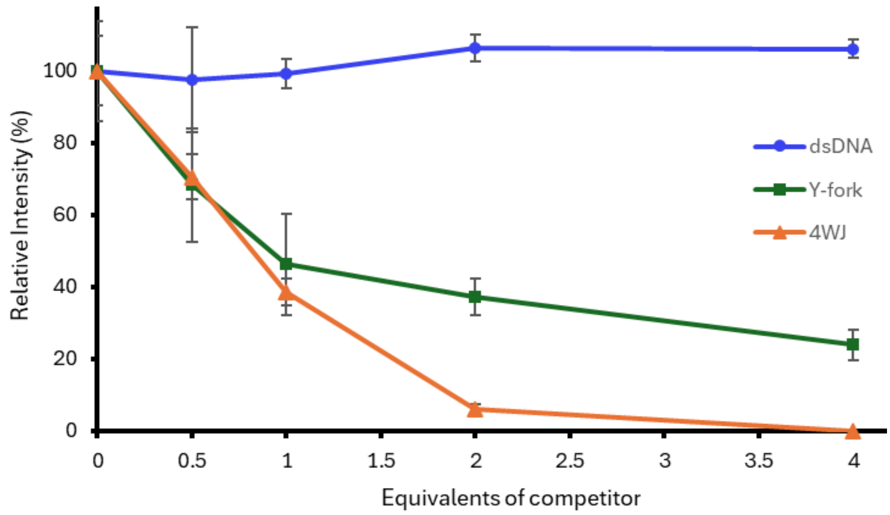


**Figure S13.** PAGE competition curves for the Pt-BIMA metallo-cage. The gel bands were measured and the relative FAM–3WJ intensity is plotted as a function of the equivalent of other DNA competitors. These curves show how Pt-BIMA has a strong preference for the 4WJ (and Y-forks to a lesser extent), in contrast to the curves in Fig. 3B, which show that the Pt helicate does not exhibit this same behaviour.

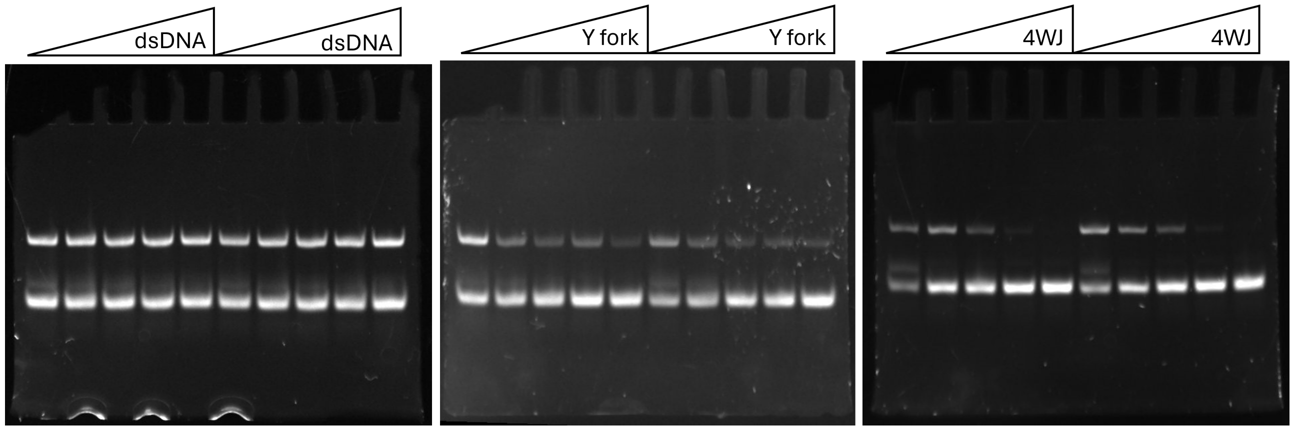


**Figure S14.** Representative gels for the Pt-BIMA PAGE competition experiments. Each mini gel contained two repeats. Each sample contained 2 µM FAM–3WJ and 1 equivalent Pt-BIMA with increasing concentrations of DNA competitor (0, 0.5, 1, 2 and 4 equivalents).

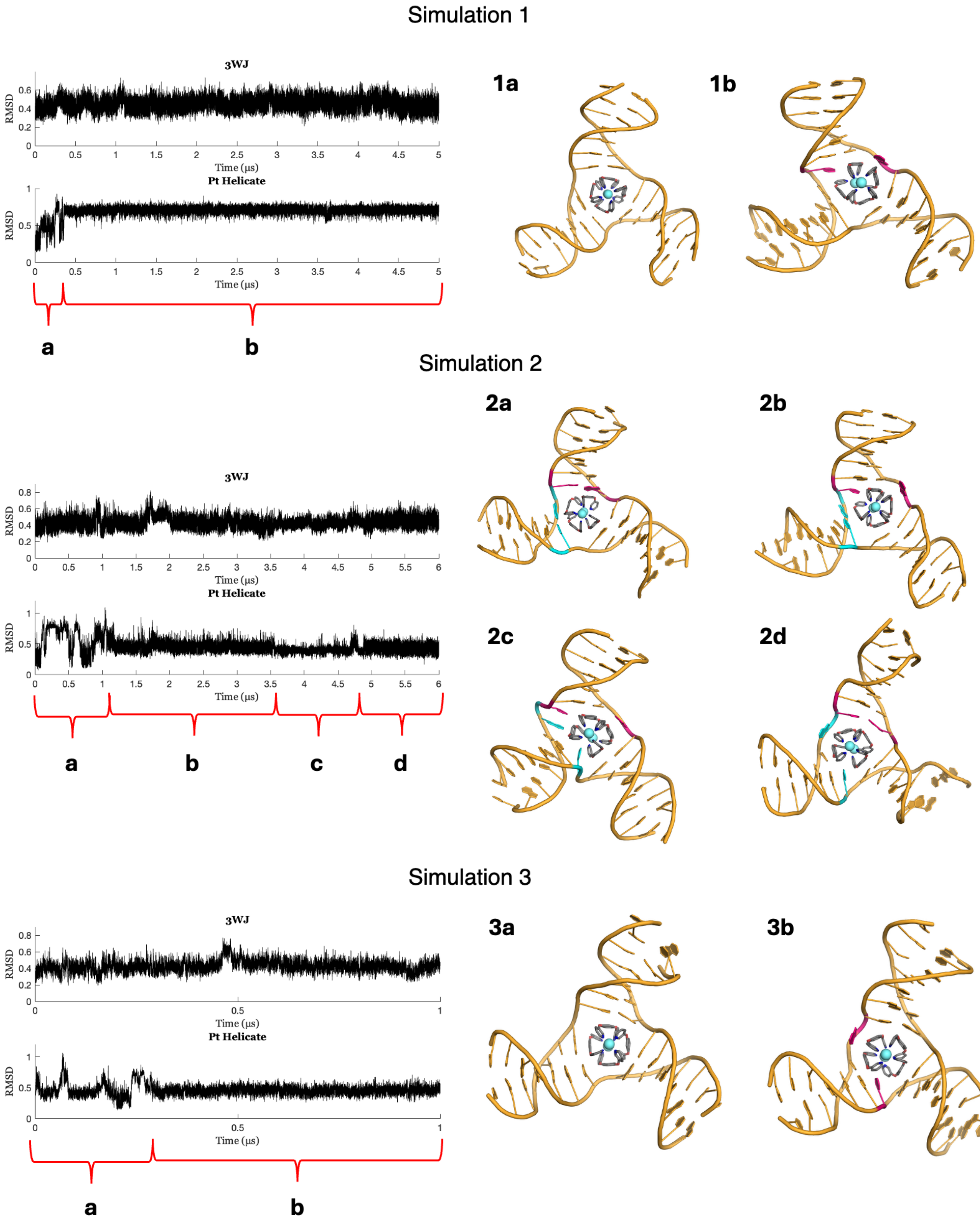


**Figure S15.** RMSD plots of the 3WJ + Pt helicate (*P* starting helicity) simulations and representative snapshots for each portion of the simulations. RMSD plots compare the 3WJ or the Pt helicate in each simulation frame to their starting coordinates. In all simulations, the compound initially rotates inside the cavity, unable to properly stack with the bases (1a, 2a, 3a) until the pink base pair opens, when it then finds its preferred binding mode (1b, 2b, 3b). In simulation 2, a second base pair opening (the cyan base pair) occurs (2c), leading to a transition into an equivalent binding mode (to 2b) at the adjacent base pair (2d).

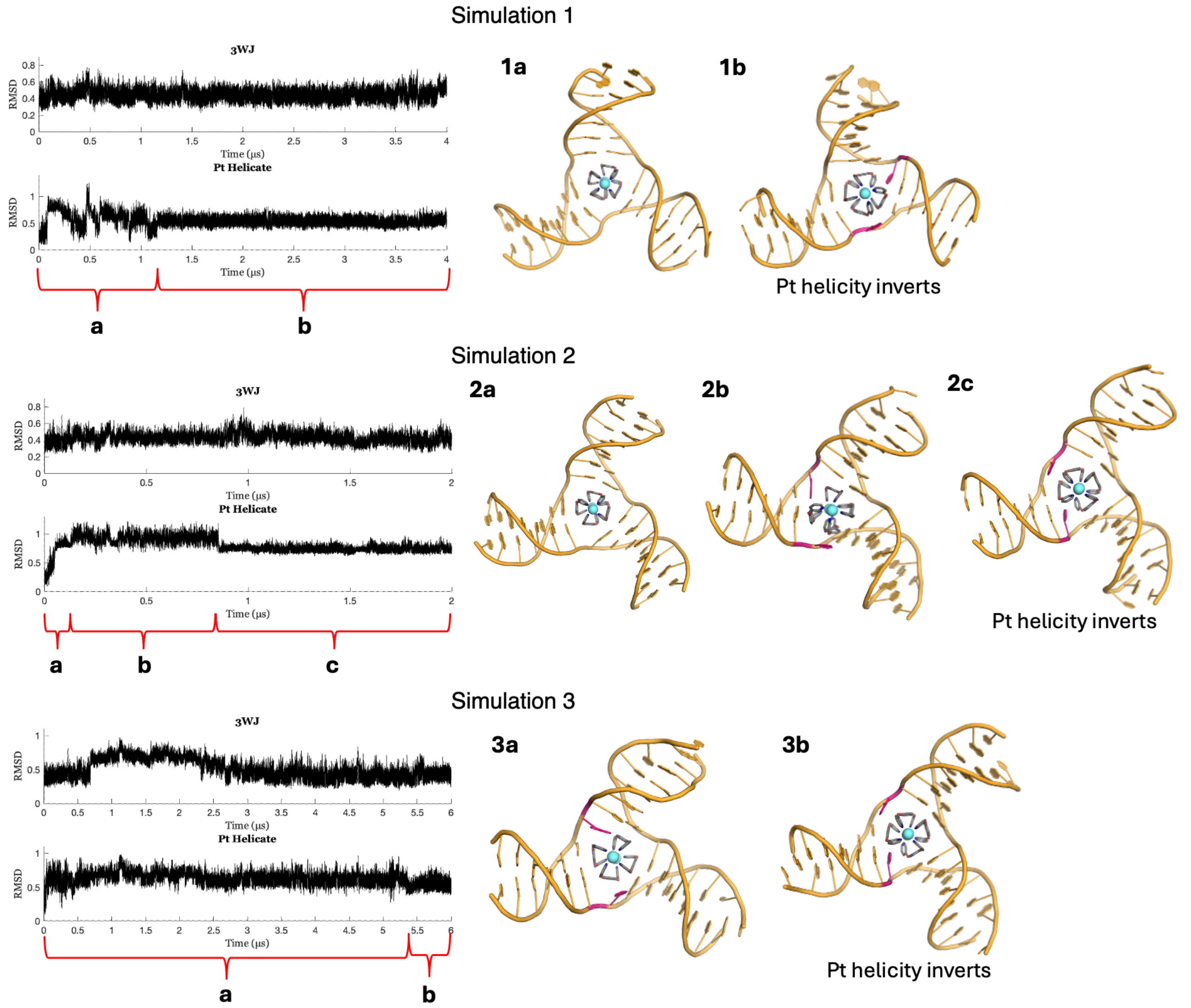


**Figure S16.** RMSD plots of the 3WJ + Pt helicate (*M* starting helicity) simulations and representative snapshots for each portion of the simulations. The RMSD plots compare the 3WJ or the Pt helicate in each frame to their starting coordinates. In simulations 1 and 2, the compound initially rotates inside the cavity, unable to properly stack with the bases (1a, 2a). In simulation 1 a base pair opens, and at the same time, the helicity of the compound flips to *P* (1b). In simulation 2, these two events happen separately: first the base pair opens, and the helicate adopts a metastable position with its *M* helicity (2b), then eventually the helicity flips and the preferred binding mode is adopted (2c). Simulation 3 captured rapid fraying of a base pair leading to a metastable binding mode for the compound (3a) – the RMSD plot indicates that there is much conformational variation for the 3WJ here. Eventually, one of those frayed bases moves back in to stack with the DNA arm and the helicity of the compound flips, allowing the compound to adopt the preferred binding mode as seen in all other simulations (3b).

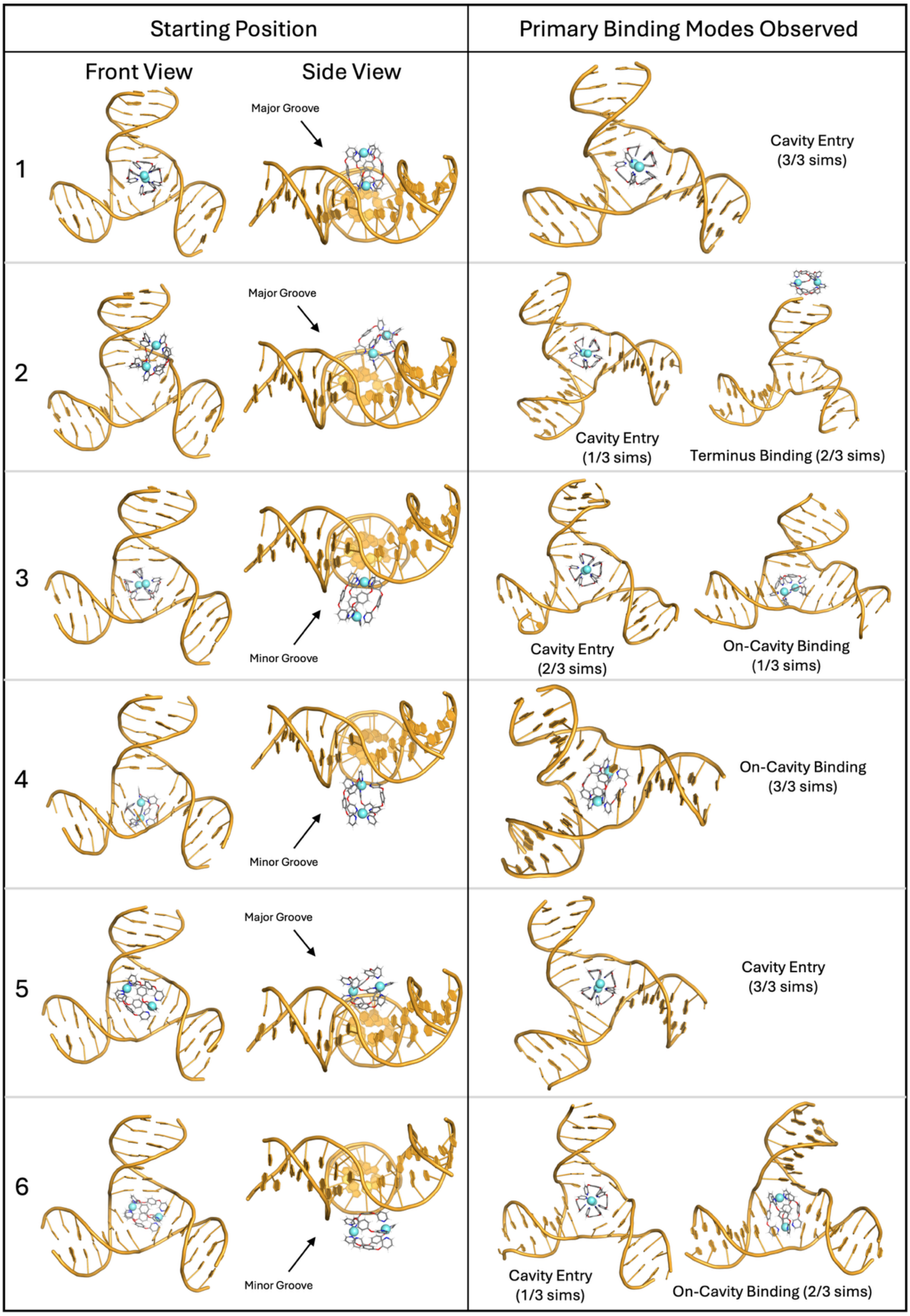


**Figure S17.** Initial positions for simulations where the Pt helicate is placed outside of the junction cavity and MD snapshots illustrating the major observed binding events.

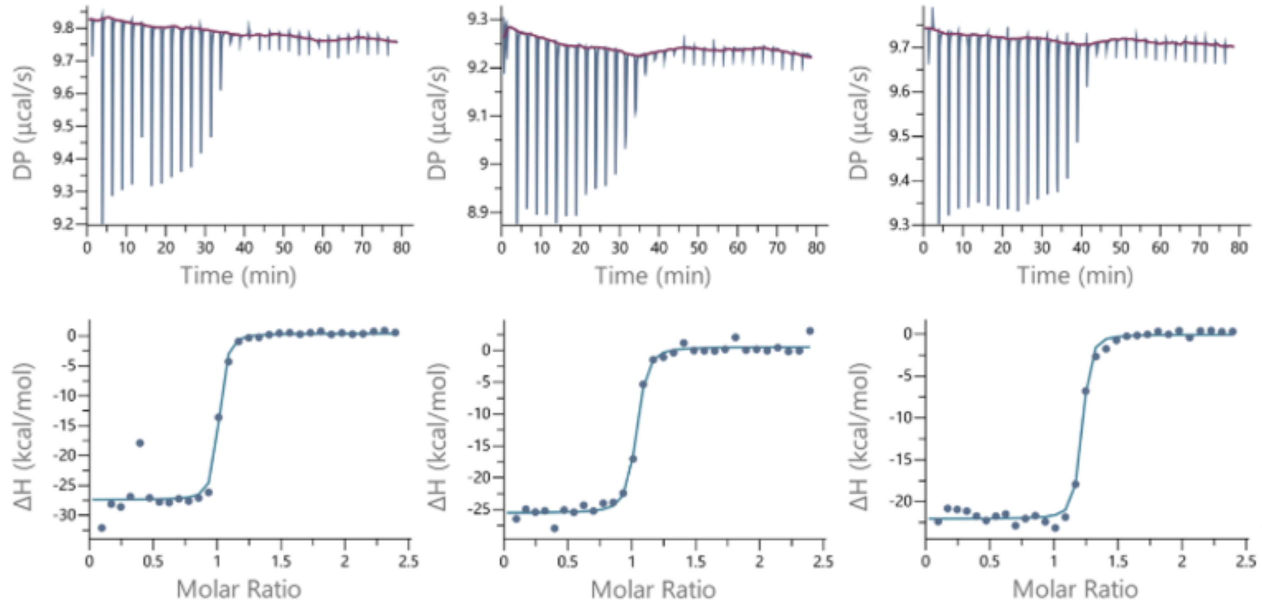


**Figure S18.** ITC data for the binding event of $Pt_2(L_1)_4(NO_3)_4$ with 3WJ-$T_6$ in buffer (10 mM sodium cacodylate, 100 mM NaCl, pH 7.4).

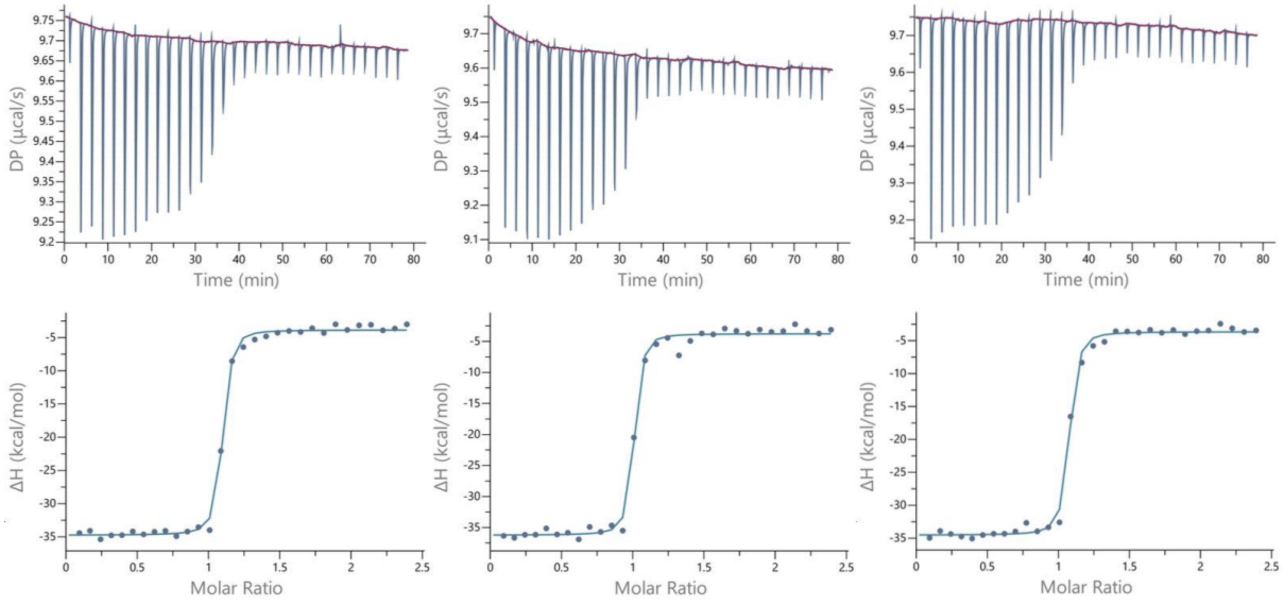


**Figure S19.** ITC data for the binding event of the Ni cylinder with 3WJ-$T_6$ in buffer (10 mM sodium cacodylate, 100 mM NaCl, pH 7.4).

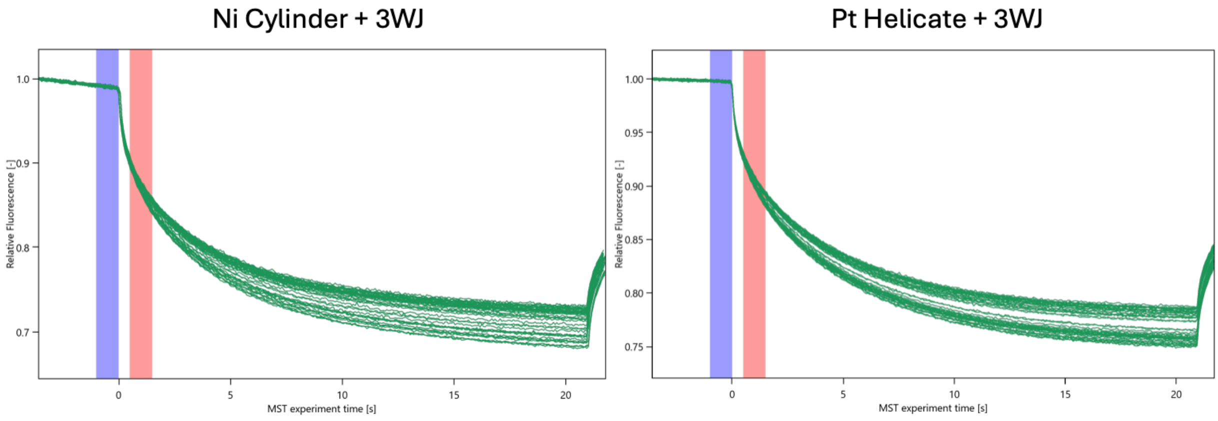


**Figure S20.** MST traces for Ni cylinder + 3WJ (left) and Pt helicate + 3WJ (right) (all repeats). The red shading represents the time range in which the average fluorescence intensity was measured to obtain the binding curve (1.5–2.5 s after IR irradiation) and the blue shading represents the initial fluorescence, to which the measurements are normalised.

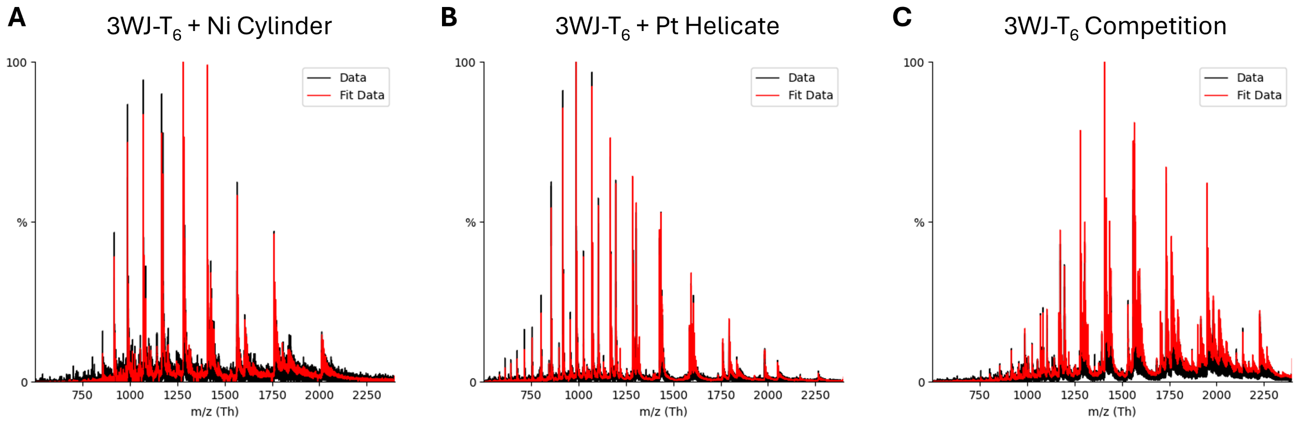


**Figure S21.** Raw ESI-MS spectra (black) and fit for deconvolution (red) for A) 3WJ-$T_6$ + Ni cylinder; B) 3WJ-$T_6$ + Pt helicate; and C) the 3WJ-$T_6$ competition study containing 1 equiv. each of Ni cylinder and Pt helicate.

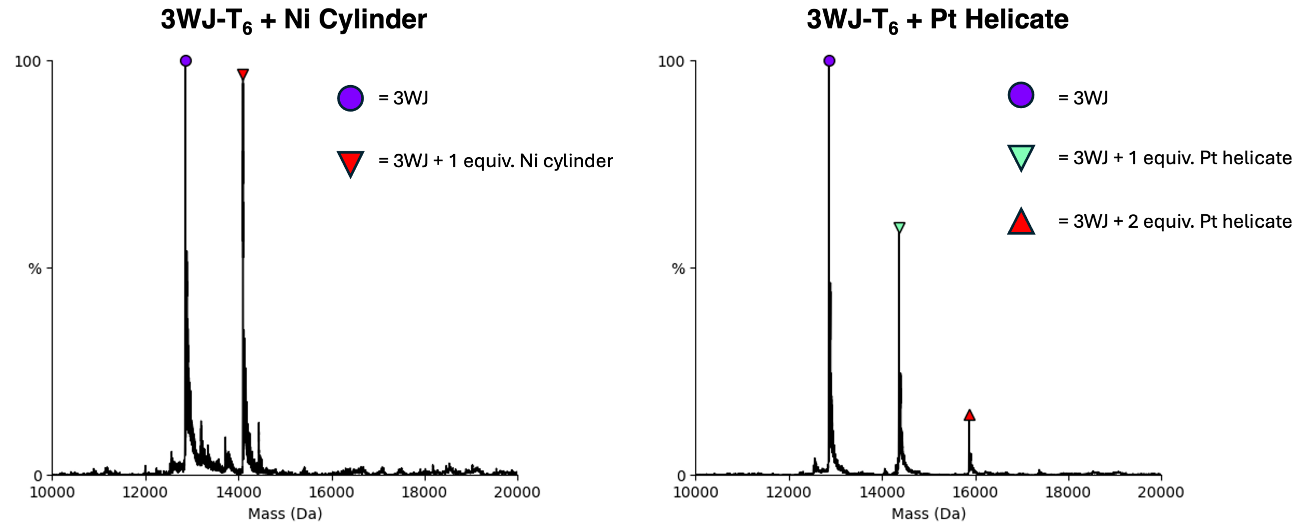


**Figure S22.** Deconvoluted mass spectra of 3WJ-$T_6$ + (left) 1 equiv. Ni Cylinder and (right) 1 equiv. Pt helicate (10 mM Ammonium acetate, 10% MeOH, pH 7.2)

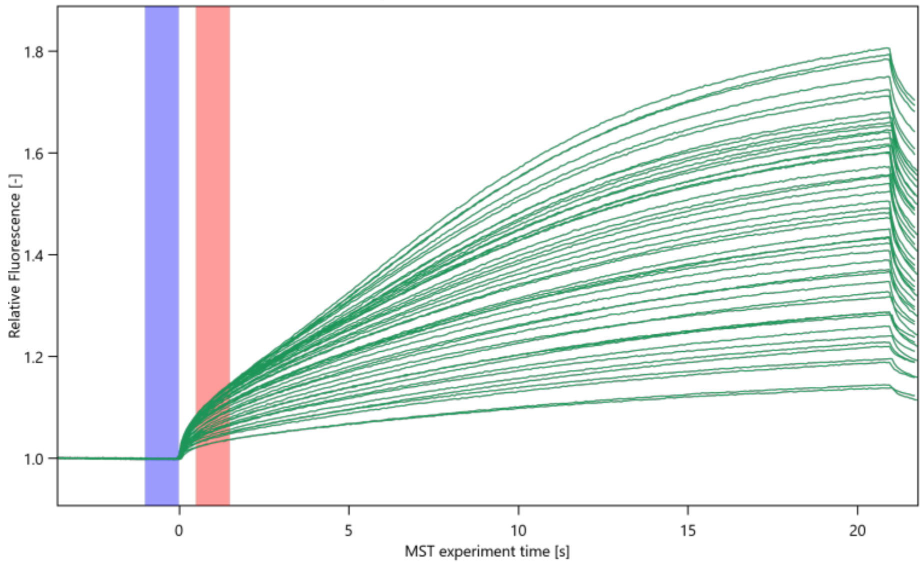


**Figure S23.** MST traces for Pt helicate + 4WJ (all repeats). The red shading represents the time range in which the average fluorescence intensity was measured to obtain the binding curve (1.5–2.5 s after IR irradiation) and the blue shading represents the initial fluorescence, to which the measurements are normalised.

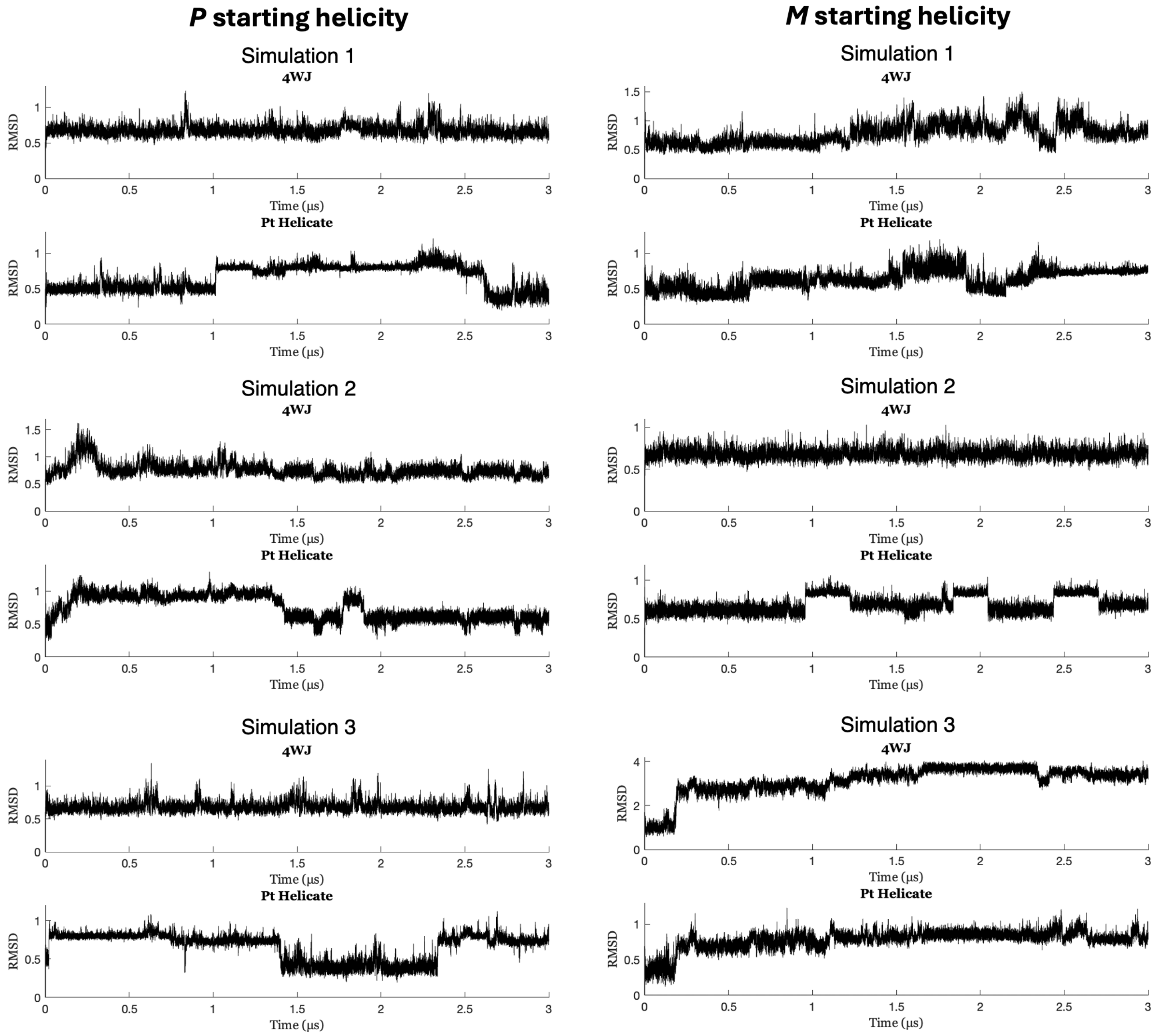


**Figure S24.** RMSD plots of the 4WJ + Pt helicate simulations and representative snapshots for each portion of the simulations. The RMSD plots compare the 4WJ or the Pt helicate in each frame to their starting coordinates. The Pt helicate plots illustrate how the compound is unable to find a stable binding mode and moves and rotates a lot inside the 4WJ cavity. The 4WJ plots also reveal a lot of conformational variation in the DNA.

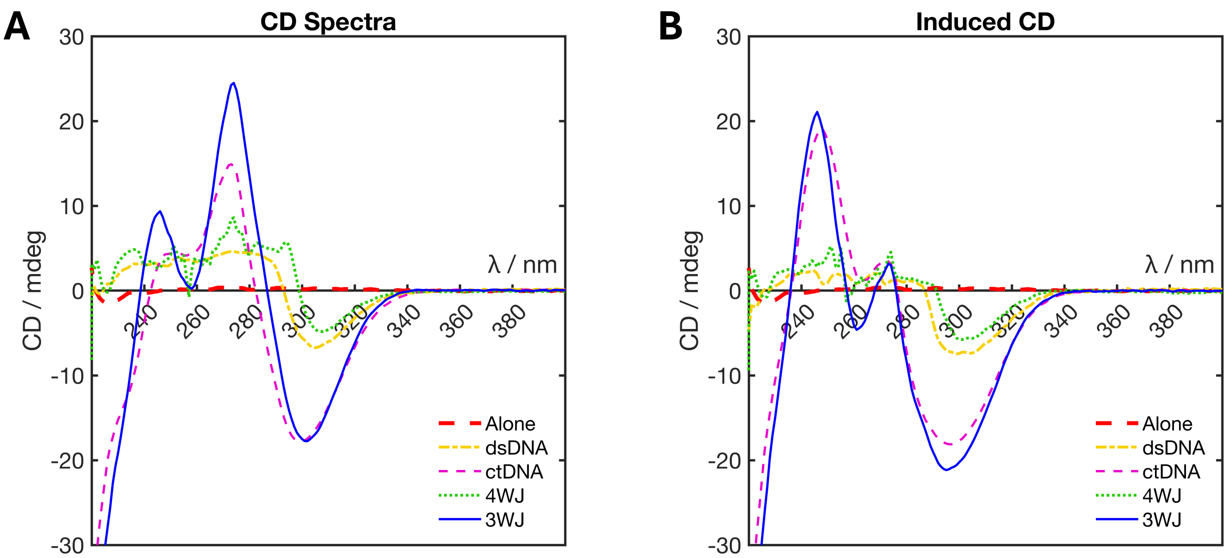


**Figure S25.** A) CD spectra and B) induced CD curves of Pt helicate alone (red) and with DS-21 (orange), ctDNA (purple), 4WJ (green) and 3WJ (blue). Induced CD curves were obtained by subtracting the CD curve of the DNA alone from the respective curves in part A.

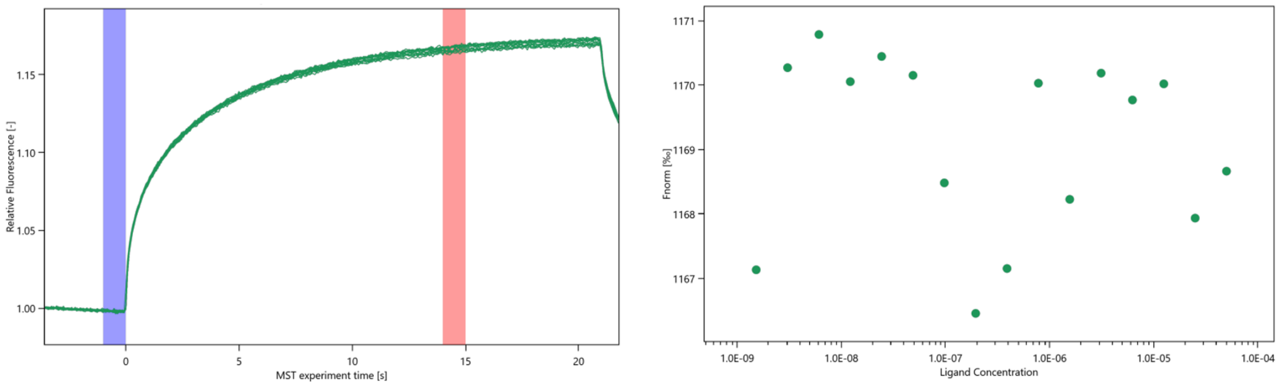


**Figure S26.** MST traces (left) and plot of average intensities (right) for the Pt helicate + DS-21. The red shading represents the time range in which the average fluorescence intensity was measured to obtain the binding curve (14–15 s after IR irradiation) and the blue shading represents the initial fluorescence, to which the measurements are normalised.

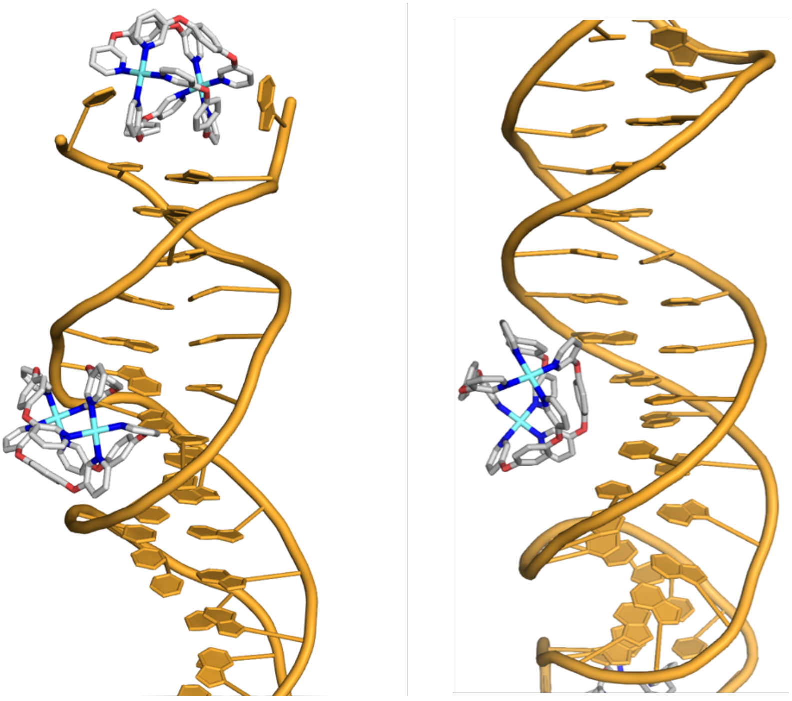

**Figure S27.** Closeup MD snapshots of the Pt helicate binding at the terminus and (transiently) in the minor and major grooves of a B-DNA 25mer.

| | Ni Cylinder | Pt Helicate |
|---|---|---|
| $K_d$ / nM (by MST) | 0.516 ± 0.67 | 12.2 ± 0.22 |
| $K_d$ / nM (by ITC) | 7.17 ± 0.51 | 9.94 ± 4.36 |
| n / sites | 1.02 ± 0.05 | 1.04 ± 0.11 |
| $\Delta H$ / kcal mol$^{-1}$ | –31.4 ± 0.9 | –25.3 ± 3.0 |
| $T\Delta S$ / kcal mol$^{-1}$ | –20.3 ± 0.9 | –14.3 ± 3.1 |
| $\Delta G$ / kcal mol$^{-1}$ | –11.1 ± 0.1 | –11.0 ± 0.3 |

**Table S1.** Thermodynamic parameters for the binding interactions of the Ni cylinder and Pt helicate with 3WJ-$T_6$, measured by MST (row 1) or ITC (rows 2–6) (10 mM sodium cacodylate, 100 mM NaCl, pH 7.4, 25 °C). Each titration was repeated 3 times.

(12) Case, D. A.; Aktulga, H. M.; Belfon, K.; Cerutti, D. S.; Cisneros, G. A.; Cruzeiro, V. W. D.; Forouzesh, N.; Giese, T. J.; Götz, A. W.; Gohlke, H.; Izadi, S.; Kasavajhala, K.; Kaymak, M. C.; King, E.; Kurtzman, T.; Lee, T.-S.; Li, P.; Liu, J.; Luchko, T.; Luo, R.; Manathunga, M.; Machado, M. R.; Nguyen, H. M.; O'Hearn, K. A.; Onufriev, A. V.; Pan, F.; Pantano, S.; Qi, R.; Rahnamoun, A.; Risheh, A.; Schott-Verdugo, S.; Shajan, A.; Swails, J.; Wang, J.; Wei, H.; Wu, X.; Wu, Y.; Zhang,

S.; Zhao, S.; Zhu, Q.; Cheatham, T. E., III; Roe, D. R.; Roitberg, A.; Simmerling, C.; York, D. M.; Nagan, M. C.; Merz, K. M., Jr. AmberTools. *J. Chem. Inf. Model.* **2023**, *63* (20), 6183–6191. DOI: 10.1021/acs.jcim.3c01153

(20) The PyMOL Molecular Graphics System, Version 2.8, Schrödinger, LLC.
